# Learning to read increases the informativeness of distributed ventral temporal responses

**DOI:** 10.1101/257055

**Authors:** Marisa Nordt, Jesse Gomez, Vaidehi Natu, Brianna Jeska, Michael Barnett, Kalanit Grill-Spector

**Affiliations:** Department of Developmental Neuropsychology, Ruhr-Universität Bochum, 44801 Bochum, Germany.; Neurosciences Program, Stanford University School of Medicine, Stanford, CA 94305, USA.; Psychology Department, Stanford University, Stanford, CA 94305, USA.; Stanford Neurosciences Institute, Stanford University, Stanford, CA 94305, USA.

**Author notes:** <u>Corresponding author:</u> Kalanit Grill-Spector, Jordan Hall, Bldg. 420, Department of Psychology, Stanford Neurosciences Institute, Stanford, California 94305-2130.

## Abstract

Becoming a proficient reader requires substantial learning over many years. However, it is unknown how learning to read affects development of distributed visual representations across human ventral temporal cortex (VTC). Using fMRI and a data-driven, computational approach, we quantified the development of distributed VTC responses to characters (pseudowords and numbers) vs. other domains in children, preteens, and adults. Results reveal anatomical- and hemisphere-specific development. With development, distributed responses to words and characters became more distinctive and informative in lateral but not medial VTC, and in the left but not right hemisphere. While development of voxels with both positive (that is, word-selective) and negative preference to words affected distributed information, only development of word-selective voxels predicted reading ability. These data show that developmental increases in informativeness of distributed left lateral VTC responses enable proficient reading and have important implications for both developmental theories and for elucidating neural mechanisms of reading disabilities.

## Introduction

Reading is a unique human ability that is learned. Each year, over a quarter of a billion children across the globe attend primary school (grades 1-6) whose central mission is literacy instruction. Reading entails visual processing of letters and words and associating these visual inputs with sounds and language. Thus, reading involves a network of brain regions involved in vision, audition, and language. As such, prior research has examined how learning to read affects white matter tracts of the reading network (Ben-Shachar, Dougherty, & Wandell, 2007; Carreiras et al., 2009; Schlaggar & McCandliss, 2007; Wandell, Rauschecker, & Yeatman, 2012; Yeatman, Dougherty, Ben-Shachar, & Wandell, 2012) and the emergence of the visual word form area (VWFA, Ben-Shachar et al., 2011; Cantlon et al., 2011). The VWFA (Cohen et al., 2000; Dehaene, Le Clec’H, Poline, Le Bihan, & Cohen, 2002) is located within ventral temporal cortex (VTC), an anatomical expanse that processes high-level visual information (Cohen et al., 2000; Dehaene et al., 2002; Grill-Spector & Weiner, 2014; Hannagan, Amedi, Cohen, Dehaene-Lambertz, & Dehaene, 2015; Rauschecker et al., 2011). A separate body of research has shown that distributed responses across VTC have a characteristic pattern that represents the category of the visual input (Carlson, Tovar, Alink, & Kriegeskorte, 2013; Golarai, Liberman, & Grill-Spector, 2017; Haxby et al., 2001; Kriegeskorte, 2008). Indeed, independent classifiers can decode from distributed VTC responses the category of the viewed stimulus (Cox & Savoy, 2003; Grill-Spector & Weiner, 2014; Weiner & Grill-Spector, 2010). However, it is unknown how learning to read during childhood affects the development of distributed representations of words and characters across VTC and if cortical development has behavioral ramifications.

To address these gaps in knowledge, we examined three questions in school-aged children and young adults.

1. *If and how does learning to read affect distributed responses across the VTC?* We considered two main developmental hypotheses. One hypothesis predicts that learning to read leads to the emergence of new distributed representations of words across VTC. This hypothesis predicts that distributed VTC responses to words and characters will become more distinct and informative from childhood to adulthood. A second hypothesis predicts that distributed representations for words and characters in VTC may not be different across children and adults because both groups see characters in their natural environment. Thus, by the age of 5, distributed VTC representations to characters may be fully developed, as has been reported for other domains such as places and objects (Golarai, Liberman, Yoon, & Grill-Spector, 2010; Golarai et al., 2017).
2. *Is there anatomical specificity to the development of distributed VTC responses to words and characters?* The theory of object form topography (Haxby et al., 2001) suggests that visual category information is obtained by distributed responses across the entire VTC (Carlson et al., 2013; Connolly et al., 2012; Cox & Savoy, 2003; Haxby et al., 2001; Kriegeskorte, 2008). This theory predicts no anatomical specificity within VTC to the development of distributed responses. In contrast, eccentricity bias theory (Hasson, Levy, Behrmann, Hendler, & Malach, 2002; Levy, Hasson, Avidan, Hendler, & Malach, 2001; Malach, Levy, & Hasson, 2002), suggests that reading requires fine-grain visual acuity afforded by foveal vision. This theory predicts that foveation on words during reading will lead to the development of word representations in cortical regions with a pre-existing foveal bias (higher responses to central than peripheral stimuli). In both children and adults regions lateral to the mid-fusiform sulcus (MFS, Weiner et al., 2014) are foveally-biased (Hasson et al., 2002; Levy et al., 2001; Weiner et al., 2014) and regions medial to the MFS are peripherally-biased (Hasson et al., 2002; Levy et al., 2001; Weiner et al., 2014). Thus, eccentricity bias predicts that learning to read will lead to development of distributed responses in foveally-biased lateral VTC, but not peripherally-biased medial VTC. A third theory of domain specificity, suggests that visual processing of words is accomplished by a specific region, the VWFA, as (i) it responds significantly more strongly to characters than other stimuli (Dehaene et al., 2002), (ii) it is causally involved in processing words (Gaillard et al., 2006), and (iii) it shows developmental increases in response amplitude to letters and (pseudo)words (Ben-Shachar et al., 2011; Cantlon et al., 2011). Thus, domain-specificity predicts that learning to read will induce an even more anatomically-specific development of distributed responses restricted to just word-selective voxels rather than the entire lateral VTC.
3. *Does development of distributed VTC responses have behavioral ramifications?* One possibility is that the development of distributed VTC responses improves reading ability, predicting a positive correlation between reading ability and information in distributed VTC responses to words and characters. If such a correlation exists, it would be critical to determine if anatomically-specific compartments of VTC predict reading ability. Alternatively, development of reading ability may depend on white matter connections between VTC with downstream areas (Gullick & Booth, 2015; Takeuchi et al., 2016; Yeatman et al., 2012) rather than distributed VTC responses. This alternative predicts no relationship between reading ability and development of distributed VTC responses.

To address these questions, we conducted fMRI in three age groups: 12 children (ages, 5-9 years; 11 female), 13 preteens (ages, 10-12 years; 6 female), and 26 adults (ages, 22-28 years; 10 female). During scanning, participants viewed images of characters (pseudowords and uncommon numbers) and items from four other domains, each consisting of two categories (Figure 1a). Participants viewed both words and numbers to allow distinguishing if development is general to the domain of characters or is specific to words. We used pseudowords, which are enunciable but lack meaning, for two reasons: (i) to control for age-related differences in semantic knowledge and (ii) to control familiarity across domains, as items from other domains were also unfamiliar.

**Figure 1.**
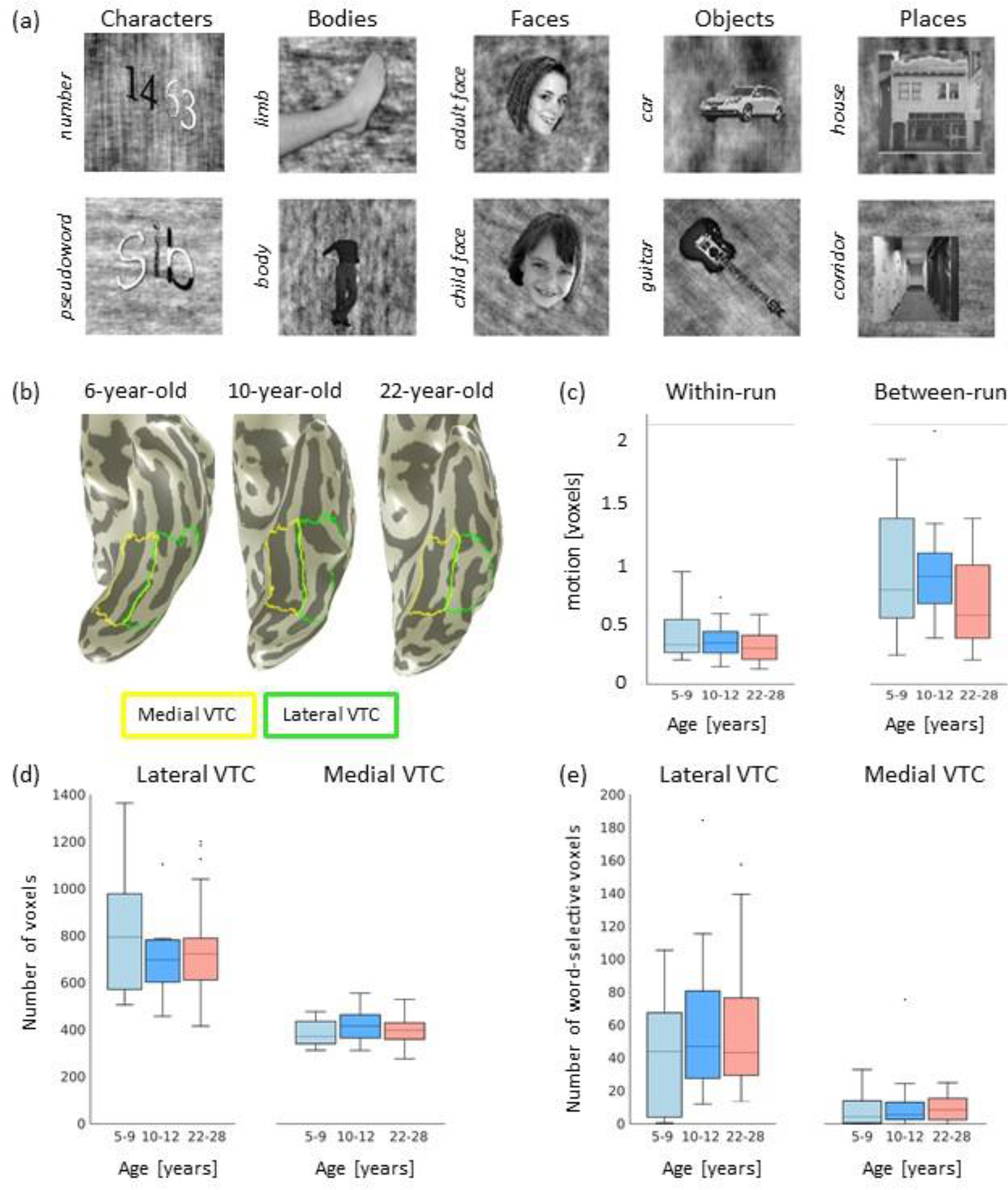
Stimuli, anatomical ROI definitions, and between age-group controls. (a) Examples of stimuli corresponding to the five domains (in columns) with two categories per domain (rows), (b) Examples of lateral and medial ventral temporal cortex (VTC) on the inflated cortical surface of a representative 6-year-old child (left), 10-year-old child (middle), and 22-year-old adult (right). *Yellow outline:* medial VTC; *green outline:* lateral VTC. (c) Boxplotsshowing within-run motion (left) and between-run motion (right) for each of the three age groups. *Light blue:* children (5-9 year old, n = 12); *blue:* preteen (10-12 year old, n = 13); orange: adults (22-26 year old, n = 26). There were no significant differences across age groups, (d) Boxplot of the number of functional voxels in lateral (left) and medial (right) VTC averaged across hemispheres. Each functional voxel is 2.4 mm on a side. Same subjects as in (c). (e) Boxplot showing the number of word-selective voxels in lateral (left) and medial (right) VTC averaged across hemispheres. Number of subjects and voxel size same as (d). In (c)-(e) the box indicates 25%-75% quartiles of the data; *Horizontal tines in the box plots:* median; *whiskers:* data range excluding outliers, encompassing 99.3% of the data; *black dots:* outliers, values that are more than 1.5 times the interquartile range away from the top or bottom of the box.

To test developmental hypotheses, we measured distributed responses to items from each category in each of the medial and lateral VTC compartments and examined: (i) if there are age-related differences in the information and discriminability of distributed VTC responses to words and characters, (ii) if development of distributed responses shows anatomical specificity, and (iii) if development of distributed VTC response predicts reading ability.

## Results

There were no significant differences across age groups in (i) motion during fMRI (Figure 1c, *F(2,48)*=2.67, *p*=0.08), (ii) the number of voxels in anatomical partitions of VTC (Figure 1d, all *Fs*≤0.86, *ps*>0.42, analysis of variance (ANOVA) with the factor age group) and (iii) the number of word-selective voxels (Figure 1e, all *Fs*≤0.71, *ps*>0.49, ANOVAs with factor age group). These analyses show that data quality and the anatomical size of VTC is similar across age groups.

### Does information of distributed VTC responses for characters, words, and numbers develop after age 5?

We used a decoding approach to quantify if there are developmental changes either in decoding characters, words, or numbers from distributed VTC responses. Thus, we quantified using a winner-takes-all classifier three types of information in distributed VTC responses: (i) characters (pseudowords+numbers) vs. other domains (faces, places, objects, and body parts), (ii) pseudowords vs. the other nine categories (adult faces, child faces, houses, corridors, cars, guitars, bodies, limbs, Figure 1a), and (iii) numbers vs. the other nine categories.

Results show that in all age groups, VTC partitions, and hemispheres, decoding character information was significantly higher than the 20% chance level (Figure 2a, all *ps*≤0.005). Notably, decoding character information significantly increased from age 5 to adulthood (Figure 2a, main effect of age group, *F*(2,48)=5.74, *p*=0.006, 3-way rmANOVA with factors of age group, VTC partition, hemisphere).

**Figure 2.**
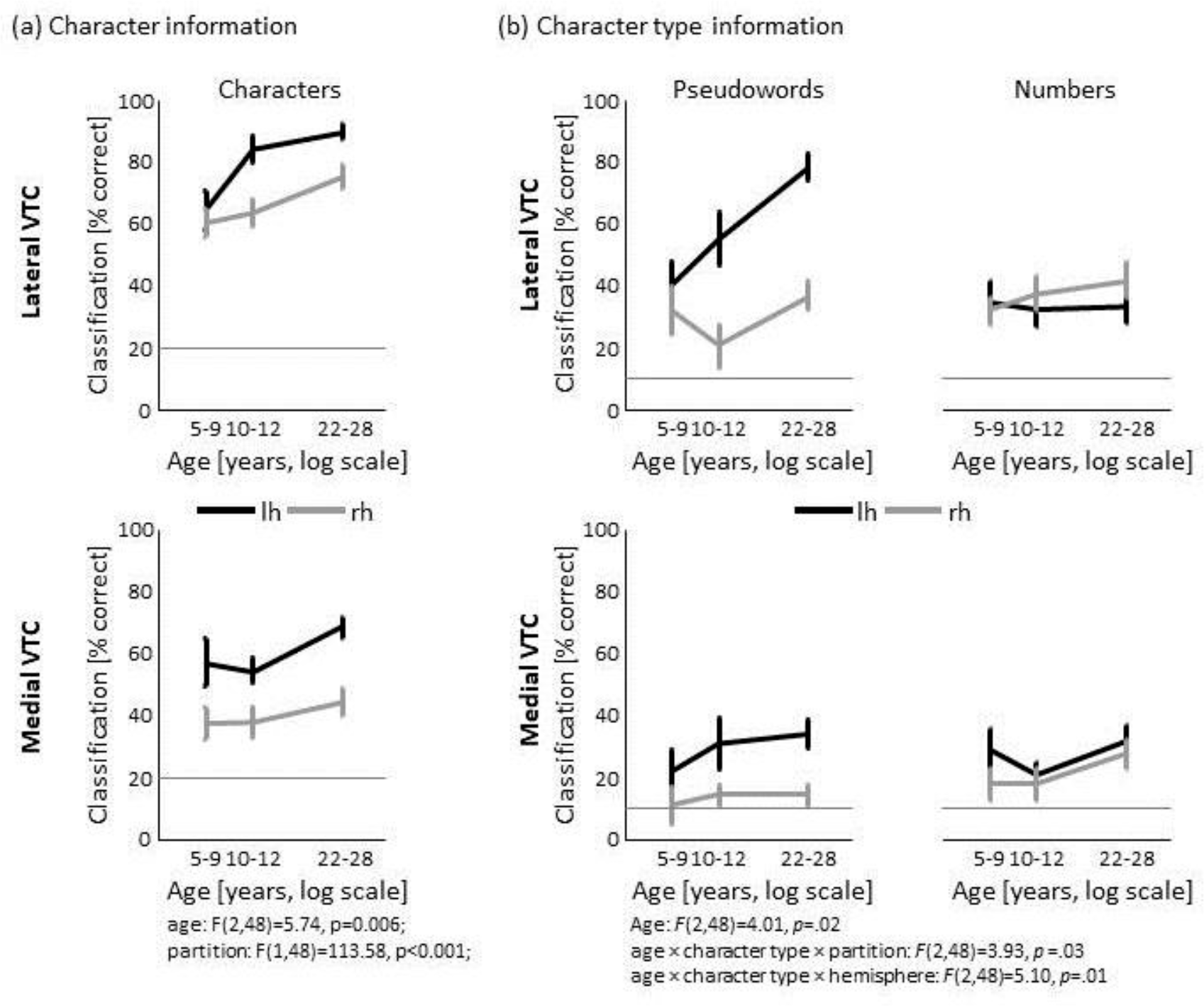
Differential development of word and number information after age 5. (a) Character classification performance in lateral VTC (top) and medial VTC (bottom) for the left (black) and right (gray) hemispheres across age groups. Classification performance was quantified with a winner-take-aii (WTA) classifier. Data show mean performance for children (5-9-year-olds, n = 12), preteens (10-12-year-olds, n = 13), and adults (22-28-year-olds, n = 26). *Error bars:* standard error of the mean (SEM). Chance level is 20% (horizontal gray line), (b) WTA classification of character type (either pseudowords or numbers) in lateral VTC (top) and medial VTC (bottom). Chance level is 10%. Same conventions as in (a). See *also* Figures S1 *and* S2.

Critically, this development was anatomically specific: character information was better decoded from multivoxel patterns (MVPs) across lateral VTC compared to medial VTC (main effect of VTC partition, *F*(1,48)=113.58,*p*<0.001), and also better decoded from left than right hemisphere MVPs (main effect of hemisphere, *F*(1,48)=40.17, *p*<0.001, no significant interactions, all *ps*>0.066). The largest development was observed in left lateral VTC in which decoding of character information improved on average by 25% from age 5 to 25. Specifically, decoding characters vs. other stimuli from left lateral VTC yielded an accuracy of 65%±6% (mean±SEM) in children, but reached a 90%±3% accuracy in adults (significantly higher than 5-9 year-olds, post-hoc *t*-test, *t(36)=4.2*, *p*<0.001, Figure 2a).

In contrast to the development of character information, we did not find a significant development of information for the domains of bodies, objects, and places in VTC (Figure S1a). However, we found a significant development of face information in VTC, consistent with prior research (Golarai et al., 2017). Critically, decoding of domain information was not due to low-level features, as decoding domain information from VTC was significantly higher than from V1 (Figure S1b, main effect of **ROI**, *F*(2,72)=142.1,*p*<0.001, rmANOVA with the factors group and ROI (lateral VTC, medial VTC, V1) in a subset of subjects with V1 defined retinotopically (5-9 year-olds, n=8, 10-12 year-olds, n=12, 22-28 year-olds, n=19). Together, these analyses suggest that information about some domains is adult-like in VTC in 5-9 year olds, even as character information continues to develop.

Surprisingly, analysis of pseudoword and number decoding revealed a differential development of word and number information across VTC partitions (Figure 2b, age group × character type × VTC partition interaction, *F*(2,48)=3.93, *p*= 0.03, 4-way rmANOVA with factors of age group, character type, partition, hemisphere), as well as a differential development of word and number information across hemispheres (age group × character type × hemisphere interaction, *F*(2,48)=5.10,*p*=0.01).

That is, pseudoword information specifically developed in the left lateral VTC, even as decoding pseudowords was significantly higher than chance in all age groups (all *ps*<0.003). Notably, decoding of pseudowords from left lateral VTC progressively increased from 40%±7% in 5-9 year-olds (significantly lower than adults, post-hoc *t*-test, *t*(36)=−2.95, *p*=0.006), to 55%±8% in 10-12 year-olds (significantly lower than adults, post-hoc *t*-test, *t*(37)=−2.72, *p*=0.01), to 78%±5% in 22-28 year-olds (Figure 2b **- top left).** This reflects an almost two-fold improvement in decoding word information from left lateral VTC from age 5 to adulthood. In contrast, in the right lateral VTC decoding pseudowords was not significantly different across age groups and overall lower than left lateral VTC, averaging at an accuracy of 31%±4% (not significantly different across age groups, all *ps*>0.15, Figure 2b**-top left).** Additionally, there were no significant differences across age groups in decoding pseudowords from medial VTC (all *ps*>0.19, Figure 2b**-bottom left)**, and performance was around 30% in the left hemisphere and only around 13% in the right hemisphere.

In contrast to the development of word information, there was no significant development of number information in any partition or hemisphere (no significant effects of age group or interactions with age group, all *Fs*<1.03, *ps*>0.36, 3-way rmANOVA on number classification with the factors age group, partition, hemisphere). Indeed, number decoding averaged about 30%±4% across age groups, partitions, and hemispheres (Figure 2b**-right).** Additionally, there was no significant development of information for other categories in VTC, except for corridors (Figure S2). Thus, information in VTC for **8** out of **10** categories remained stable from childhood to adulthood. This provides evidence that the increased information for pseudowords from lateral VTC responses is specific, and does not reflect a general developmental increase in category information across VTC.

Taken together, these analyses reveal two important findings. First, we find evidence for development of both character and word information in VTC. Second, the development of character information occurs across the lateral VTC in both hemispheres, but development of word information is largely restricted to left lateral VTC.

### Is the developmental increase in word and character information driven by changes in within-domain similarity or between-domain distinctiveness?

We next sought to determine if development of word and character information is due to (i) an increase in the similarity between MVPs within the character domain, (ii) decrease in the dissimilarity (increase in distinctiveness) between MVPs of characters vs. other domains or (iii) increases in both within-character-domain similarity and between-domain distinctiveness. Similarity was estimated by computing the Pearson correlation coefficient between MVPs to different items from different runs.

Results reveal three main findings. First, across age groups and VTC partitions, within-character-domain correlations between MVPs to pseudowords (w-w) and numbers (n-n) were positive (Figure 3, *ps*<0.001), indicating that MVPs generalize to other items within the category. Second, crucially, within-character-domain similarity of word MVPs (w-w) and number MVPs (n-n) systematically increased from age 5, to age 12, to adulthood (main effect of age, *F*(2,48)=4.16, *p*=0.02, 4-way rmANOVA with factors age group, hemisphere, VTC partition, character type). Third, development of the reliability of pseudoword and number MVPs was anatomically heterogeneous. Development significantly varied across hemispheres and VTC partitions (significant age group × hemisphere × VTC partition interaction, *F*(2,48)=4.21,*p*=0.02), as well as across hemispheres and character types (significant age group × character-type × hemisphere interaction, *F*(2,48)=3.52, *p*<0.04, Figure 3,w-w/n-n).

**Figure 3.**
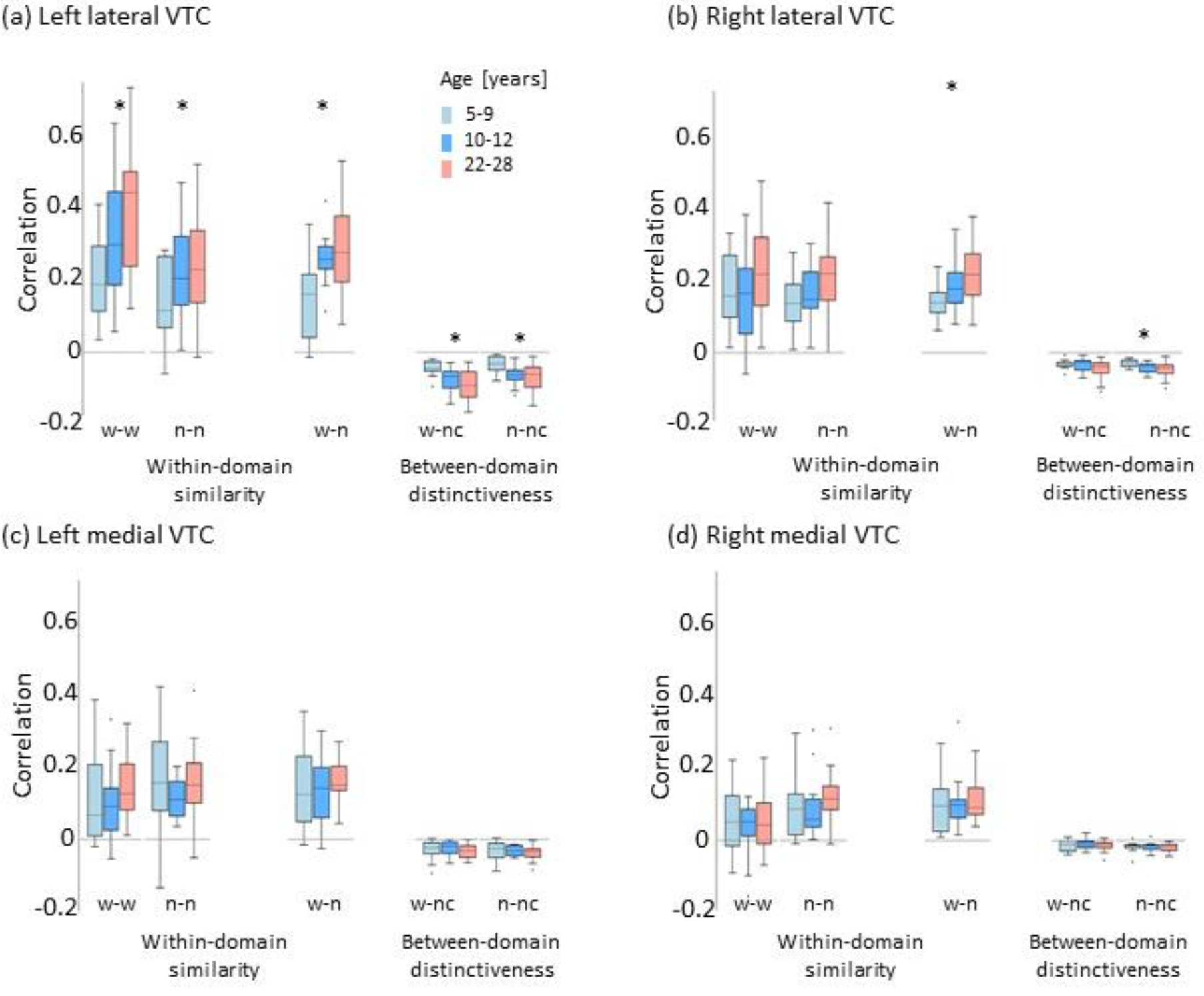
Within-domain similarity and between-domain distinctiveness of distributed responses to characters increase from age 5 to adulthood. Boxplots depict the Pearson correlation coefficient between multivoxel patterns (MVPs) of distributed responses within the domain of characters (w-w: pseudowords-pseudowords; n-n: numbers-numbers; w-n: pseudowords-numbers) and between domains (w-nc: pseudowords-non characters (all stimuli except words and numbers); n-nc: numbers-non characters, (a) Left lateral VTC; all comparisons showed a significant main effect of age as indicated by the asterisk (Fs(2,48) > 3.34; ps< 0.04). (b) Right lateral VTC; w-n and n-nc showed a significant effect of age (Fs(2,48) > 4.05; *ps*<0.02). (c) Left medial VTC. (d) Right medial VTC. Boxplots are colored by age; *light blue:* 5-9-year-olds, n = 12; *bright blue:* 10-12-year-olds, n = 13, *orange:* 22-28-year-olds, n = 26. In the boxplots, horizontal lines indicate the median, whiskers correspond to approximately ±2.7 standard deviations which captures 99.3% of the data, and extend to the most extreme value that is not an outlier. *Asterisks:* Significant main effect of age. See *also* Figure S3.

The largest development of within-character-domain similarity of MVPs occurred for pseudowords in left lateral VTC, as compared to pseudowords or numbers in medial VTC or the right hemisphere. Indeed, similarity between pseudoword MVPs in left lateral VTC increased from a median value of .19±.18 (median±interquartile range) in 5-9 year-olds to, 44±.26 in adults (Figure 3a, w-w). In contrast, similarity between pseudoword MVPs in left medial VTC only increased from .07±.20 to .13±.13 from age 5 to adulthood (Figure 3c, w-w), and the similarity of distributed responses to numbers in left lateral VTC, increased from a median of .12±.20 to .23±.20 from age 5 to adulthood (Figure 3a, n-n). In lateral VTC, pseudoword MVPs also became more similar to number MVPs from age 5 to adulthood (Figure 3, w-n, significant age group × VTC partition interaction, *F*(2,48)=9.33, *p*<0.001, 4-way rmANOVA).

Evaluation of between-domain correlations revealed that in all age groups words and numbers generate MVPs that were distinct from items of other domains (Figure S3 shows the 39 entire representational similarity matrix). Indeed the correlation between MVPs to words (or numbers) vs. items of other-domains were negative (Figure 3, w-nc/n-nc) and were significantly lower than the within-character-domain correlations (main effect of domain type, lateral VTC: *F*(1,48)=330.4, *p*<0.001; medial VTC: *F*(1,48)=157, *p*<0.001, rmANOVA with factors age group, hemisphere, domain type (within-domain/between-domain)).

Notably, MVPs to both words and numbers became significantly less correlated (or more dissimilar) from MVPs to other domains from age 5 to adulthood (Figure 3, w-nc/n-nc, main effect of age, *F*(2,48)=4.57, *p*<0.02, 4-way rmANOVA with factors age group, VTC partition, hemisphere, character type). This developmental decrease of between-domain similarity of character MVPs vs. other domains varied across VTC partitions (significant age group × VTC partition interaction, *F*(2,48)=9.79, *p*<0.001) as well as hemispheres (significant age group × VTC partition × hemisphere interaction, *F*(2,48)=3.96, *p*=0.026). Between-domain correlations significantly decreased in left lateral VTC, for both pseudowords and numbers (Figure 3a, w-nc/n-nc, both *Fs*≥6.07, *ps*≤0.005) and in right lateral VTC for numbers (Figure 3b, n-nc, *F*(2,48)=4.05, *p*=0.01).

Together, these analyses reveal that the increase in word and character information in lateral VTC appears to be driven by both increases in within-character-domain similarity as well as increases in dissimilarity (distinctiveness) across domains.

### Which voxels drive development of distributed responses for words and characters in lateral VTC?

We observed that the largest development of word and character information is in the left lateral VTC. As the lateral VTC contains the visual word form area (VWFA), this raises the question whether the development of the VWFA, which responds more strongly to characters and words vs. the other domains, is what drives the development of word and character information in lateral VTC. Alternatively, it may be that development of the entire lateral VTC including non-selective voxels increases word and character information.

To test these hypotheses, we examined classification performance separately for selective and non-selective voxels for pseudowords and characters within the lateral VTC, respectively. The first hypothesis predicts that information in selective voxels, but not non-selective voxels will increase across development. The second predicts that information in both types of voxels will increase. Selective voxels were defined in each subject and hemisphere as in prior studies (Golarai et al., 2010; Golarai et al., 2017; Weiner & Grill-Spector, 2010) as voxels that responded more strongly to the preferred stimulus compared to other stimuli. In our analyses, we separately considered voxels in lateral VTC that were word-selective (pseudowords>non-words, *t*>3, voxel-level), and character-selective (pseudowords+numbers> non-characters, *t*>3, voxel level). Non-selective voxels were defined as the remaining lateral VTC voxels. We then examined if the WTA classifier can (i) classify word and character information from the selective and non-selective voxels and (ii) if there is differential development of information across the two voxel types.

Results are consistent with the second hypothesis. First, significant development of word information occurred in both selective and non-selective voxels (main effect of age, *F*(2,42)=5.39, *p*=0.008, 3-way rmANOVA with factors of age group, hemisphere, voxel type (selective/non-selective), and no age group × voxel type interaction, *F*(2,42)=0.26, *p*=0.77) left hemisphere: Figure 4a; right hemisphere: Figure S4a). Second, similar results were obtained for character classification (Figure 4d, Figure S4d, main effect of age, *F*(2,47)=12.79, *p*<0.001 3-way rmANOVA with factors of age group, hemisphere, voxel type (selective/non-selective), and no age group × voxel type interaction, *F*(2,47)=2.13, *p*=0.13). However, conclusions from this analysis need to be considered with caution, as: (1) results may depend on the threshold used to define selectivity, (2) selective and non-selective voxel subsets are of vastly different set sizes (Figure 4a,d-legend), and (3) classification performance from these subsets is substantially lower than from all lateral VTC (Figure 4a,d, gray lines). Therefore, we sought to conduct a principled analysis that compares information in a systematic manner across threshold levels and percentage of lateral VTC voxels.

**Figure 4.**
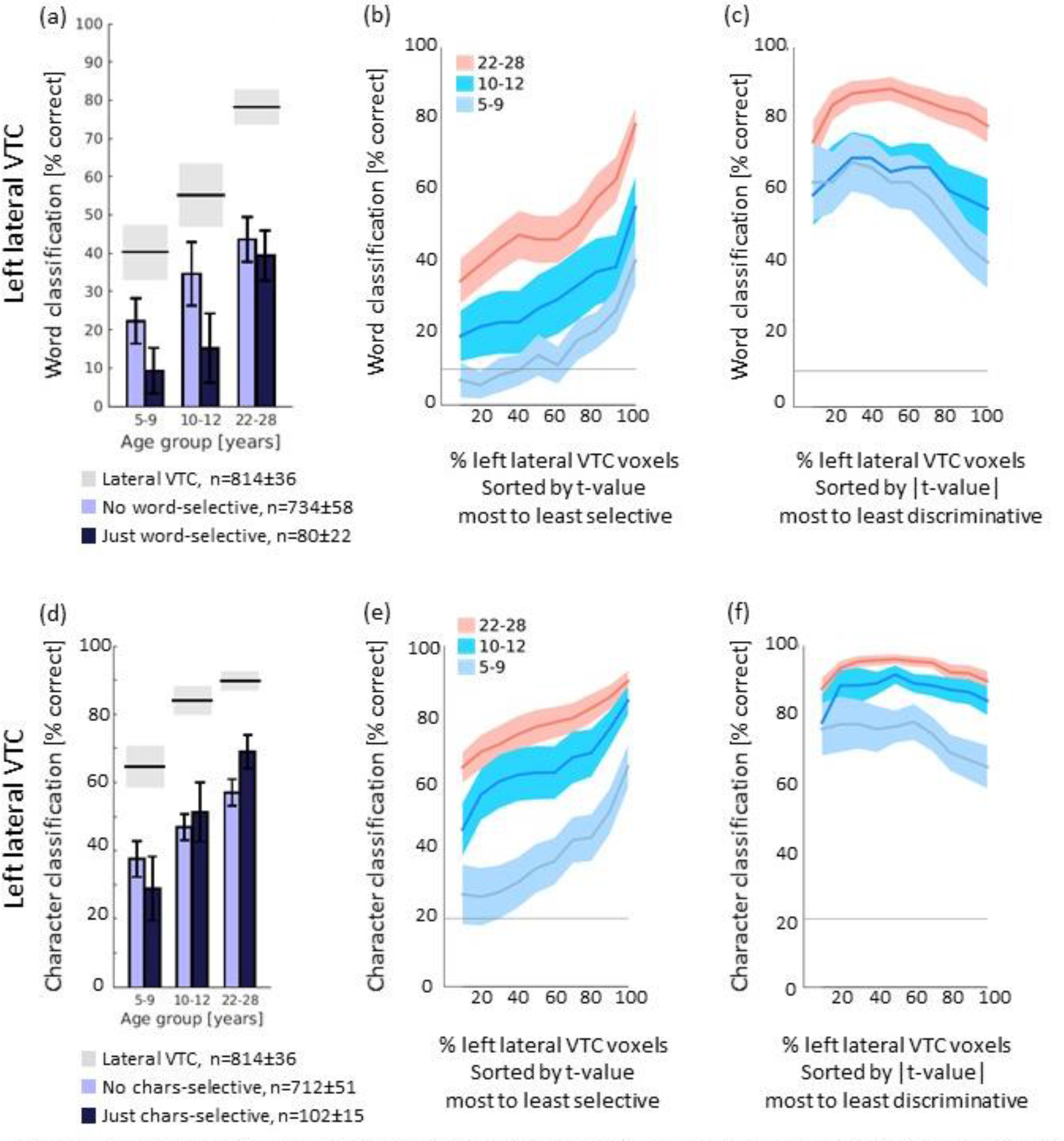
Development of word and character information in selective, non-selective, and discriminative voxels. (a) *Dark blue bars:* Mean WTA classification performance of left lateral VTC word-selective voxels (t>3). *Ugh t blue bars:* Mean WTA classification performance of left lateral VTC voxels excluding the word-selective voxels (t ≤ 3). *Horizontal lines:* mean (black) and SEM (gray) WTA classification performance from the entire left lateral VTC. *Legend:* number of voxels included in each analysis. Only subjects that had word-selective voxels are included (5-9-year-oids: *n* = 9; 10-12-year-olds: *n* = 11; 22-28-year-olds: *n* = 25). (b) Word classification performance as a function of percentage of left lateral VTCvoxels sorted from most to least word-selective (descending t-value for the contrast pseudowords>non-words). *Unes:* mean performance; *Shaded areas:* SEM. (c) Word classification performance as a function of percentage of left lateral VTC voxels sorted by descending absolute t-value (|t-value|) for the contrast pseudowords>non-words. (d.e.f) Same as a-c but for the contrast characters > non characters. In d-e only subjects that had character-selective voxels are included (5-9- year-olds: *n* = 11; 10-12-year-olds: *n* = 13; 22-28-year-olds:*n* = 26). *See also Figures S4 and S5.*

First, we tested if selective voxels drive decoding performance using a flexible approach in which we systematically varied the threshold. Thus, we sorted each subject’s lateral VTC voxels based on their selectivity to words (descending *t-value* for the contrast pseudowords>non-words) and tested classification performance as a function of number of voxels. Second, we tested if *discriminative* rather than *selective* voxels drive decoding performance. We reasoned that not only voxels with positive selectivity to words, but also voxels with negative selectivity to words may be informative. Thus, we sorted each subject’s lateral VTC voxels based on their *absolute* t-value (i.e., either positive or negative preference to words) from the greatest to least in magnitude.

Analysis by word selectivity revealed that in all age groups, word classification performance gradually increased as more voxels with progressively lower word-selectivity were added. Additionally, at all thresholds, classification in adults was higher than in children. Performance in both hemispheres was maximal when all lateral VTC voxels were included (Figure 4b, Figure S4b). Maximal decoding of pseudowords was significantly higher in adults compared to children (Figure 4b, Figure S4b, main effect of age group, *F*(2,48)=5.34, *p*=0.008, 2-way rmANOVA with factors of age group, hemisphere on estimated maximum decoding, voxels sorted by selectivity) in the left (Figure 4b), but not in the right hemisphere (Figure S4b, age group × hemisphere interaction, *F*(2,48)=6.55, *p*=0.003).

Analysis by discriminability revealed that word classification from discriminative voxels yielded maximal performance using just ~35% of lateral VTC voxels (Figure 4c, Figure S4c). Across age groups and hemispheres performance plateaued for a range of 30-60% of voxels. Including additional voxels reduced performance.

Surprisingly, pseudoword classification from the entire lateral VTC was substantially lower than the maximal classification from the discriminative subset of voxels (all *ps*≤0.005, Figure 4c, Figure S4c). For example, in 5-9 year olds, maximal classification using 36%±31% of discriminative voxels was 77%±6%, but performance dropped to 40%±7% using the entire left lateral VTC. The difference was even more pronounced in the right hemisphere: in 5-9 year olds maximal classification using 36%±21% of discriminative voxels was 64%±8%, but performance dropped to 32%± 8% for the entire right lateral VTC. The threshold of defining discriminative voxels (absolute t-value corresponding to the percentage of voxels that yielded maximal classification) did not differ significantly across age groups (no significant effect of age group, *F*(2,44)=0.15, *p*=0.86, 2-way rmANOVA on threshold absolute t-values with factors age group, hemisphere).

Critically, classification of word information from discriminative voxels developed: decoding was significantly higher in adults compared to children (Figure 4c, Figure S4c main effect of age, *F*(2,48)=7.46, *p*=0.002, 2-way rmANOVA on maximum word classification with factors age group, hemisphere). However, there was no significant development of word information in the remainder of lateral VTC voxels after the discriminative voxels that achieved highest classification were removed (no significant effect of age group, *F*(2,44)=0.48, *p*=0.62, rmANOVA with factor age group, hemisphere on non-discriminative voxels). Maximum classification from discriminative voxels was higher in adults compared to both 5-9 and 10-12 year-old children in both hemispheres (post-hoc *t*-tests, all *p*<0.02). The number of discriminative voxels yielding maximal word classification differed across hemispheres in adults, but not in children (age group × hemisphere interaction, *F*(2,37)=5.05, *p*=0.01, 2-way rmANOVA). In adults, highest classification was achieved using 51%±29 of discriminative left lateral VTC voxels vs. 24%±17% of right lateral VTC voxels.

We observed similar patterns of results when varying the percentage of character-selective (Figure 4e, Figure S4e) and character-discriminative voxels from lateral VTC (Figure 4f, Figure S4f). In contrast, there were no significant differences across age groups in decoding number information from lateral VTC using either number-selective or number-discriminative voxels (all *Fs*<1.5, *ps*>0.23, Figure S5).

These analyses lead to a surprising insight: development of word information results from developmental increases in the distinctiveness of distributed responses to pseudowords as compared to other stimuli in lateral VTC. This suggests that both voxels with the strongest positive preference to pseudowords (and characters) and those with the most negative preference contribute to word information. However, voxels with no preference (either positive or negative to pseudowords) do not contribute to classification and in fact adding many of them decreases information.

### Does word and character information in lateral VTC predict reading ability?

We reasoned that if VTC development affects reading ability, participants with more informative representations (i.e., those whose VTC produced better classification) would also read better. Thus, we examined if there is a correlation between reading performance (assessed with the Woodcock Reading Mastery Test, WRMT) outside the scanner with word and character classification performance from distributed lateral VTC responses. The WRMT tests reading accuracy by scoring how many words or pseudowords of increasing difficulty subjects can read until they make 4 consecutive errors.

Consistent with our hypothesis, WRMT performance was significantly correlated with both character classification from left lateral VTC (*r*=.60, *p*<0.001; also significant after partialling out age, *r*=.36, *p*<0.03, Figure 5a) and word classification from left lateral VTC (*r*=.58, *p*<.001; after partialling out age *r*=.22, *p*=0.19). In contrast, there was no significant correlation between WRMT performance and either word or character classification from medial VTC in either hemisphere (*rs*≤.27, *ps*>0.1). There was also no significant correlation between WRMT performance and word classification in the right lateral VTC (*r*=.19, *p*=0.45), and the correlation between WRMT performance and character classification in right lateral VTC (*r*=.33, *p*<0.05), was not significant after factoring out age (*r*=−0.08, *p*=0.64). Together, these results suggest that development of character and word information in left lateral VTC correlates with reading ability.

**Figure 5.**
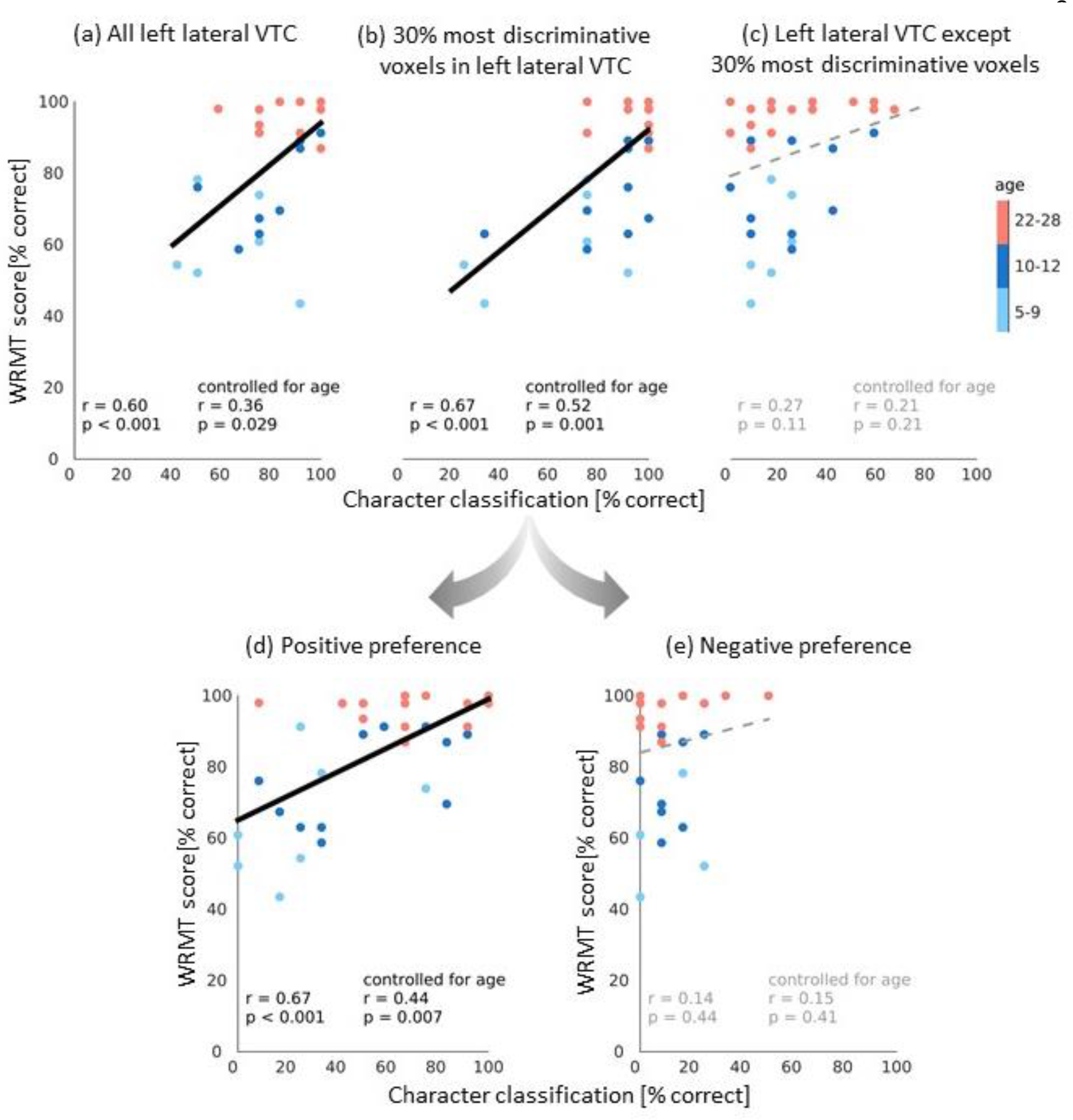
Character classification in discriminative voxels predicts reading performance. (a-c). Scatterplots of the correlation between reading ability measured by performance in the WRMT and character classification from distributed left VTC responses. Graphs differ in the brain classification data, (a) All left lateral VTC voxels, (b) 30% most discriminative voxels (see Figure 4c). (c) Left lateral VTC voxels except the 30% most discriminative voxels. Voxel data from (b) was further split into two sets, those with a positive preference to words (d) and those with a negative preference to words (e). In all plots, correlations are indicated by lines and numbers in the bottom left. *Solid black lines;* significant correlations; *Dashed gray:* non significant correlations. Only participants who completed the WRMT are shown here (5-9-year-olds, n = 7; 10-12-year-olds, n = 11; 22-28-year-olds, n = 19).

As our analyses of information in VTC reveal that discriminative voxels drive the development of character and word information in left lateral VTC, we next tested if discriminative voxels better predict reading performance than the remaining lateral VTC voxels. Results reveal that reading ability was significantly correlated with classification of characters based on the top 30% discriminative voxels (*r*=.67, *p*<0.001, also significant when partialling out age *r*=.52,*p*< 0.001, Figure 5b). In contrast, there was no significant correlation between WRMT and character classification of the non-discriminative lateral VTC voxels (*r*=.27, *p*=0.11, Figure 5c). Furthermore, the former correlation was significantly higher than the latter (Fisher transform, *p*=0.03). Likewise, WRMT was correlated with word classification based on the 30% left lateral VTC voxels which were most discriminative for pseudowords (*r=* .57, *p*<0.001; after partialling out age *r*=.28, *p*=0.1), but there was no significant correlation between WRMT and the left lateral VTC excluding these discriminative voxels (*r*=.27,*p*=0.11).

Finally, we determined which of discriminative voxels, those with positive or those with negative preference, predict reading performance. Results indicate that WMRT performance was significantly correlated with classification from voxels with positive preference to characters (*r*=.67,*p*<0.001; significant after partialling out age, *r=.*44,*p*=0.007, Figure 5d), but not with those with negative preference (*r*=.14, *p*=0.44, Figure 5e). Similarly, WRMT performance was correlated with classification based on voxels with positive preference to words (*r=.*54, *p*<0.001, after partialling out age, *r*=.31, *p*=0.07), but not with those with negative preference (*r=.*25, *p*=0.15). In other words, reading ability is predicted by distributed information from voxels with positive preference (that is, are selective) to characters and words within the top 30% discriminative voxels. Critically, this relationship dissolves when these selective voxels are excluded.

In sum, these analyses reveal that development of a subset of voxels rather than the entire left lateral VTC, best correlates with reading performance. This suggests that development of reading ability is guided by neural development that is both anatomically specific (left lateral VTC) and functionally specific (discriminative voxels), and is largely driven by the word- and character-selective voxels.

## Discussion

Reading is a complex process requiring years of practice until it is mastered. The present study investigated how learning to read during childhood affects the development of distributed representations of characters and words across VTC. Our study is the first to show that distributed responses in left lateral VTC become more distinctive and informative after age 5, and crucially that this development is linked to reading ability. This conclusion is supported by 4 observations: First, character and word information in distributed VTC responses increase from age 5 to adulthood in an anatomical and hemispheric specific manner. The most prominent development occurs in left lateral VTC compared to right lateral VTC or medial VTC, bilaterally. Second, development of character information occurs even as information for the domains of bodies, objects, and places do not develop significantly, indicating a differential development of domain information in VTC (Golarai et al., 2007, 2010; Golarai et al., 2017). Third, developmental increases in information regarding words/characters are due to development of distributed responses across the subset of discriminative word/character voxels within lateral VTC. Fourth, even as information in VTC develops in both voxels with positive and negative preference to characters/words, it is the development of selective voxels with positive preference that predicts reading ability. These findings have important implications for both developmental theories and understanding the neural bases of reading disabilities, which we detail below.

Notably, the anatomical specificity of development of word/character information in distributed VTC responses is consistent with the predictions of the eccentricity bias (Malach, Levy, & Hasson, 2002) and graded hemispheric specialization theories (Behrmann & Plaut, 2015). The former is supported by data showing that word information develops in lateral VTC, which shows a foveal bias in both children and adults (Weiner et al., 2014), but not in medial VTC, which shows a peripheral bias. The latter is supported by data revealing a more prominent development of word/character information in the left than right lateral VTC. According to this theory, when children learn to read, processing of letters and words depends more on the left hemisphere because of left lateralization of language areas in other parts of the brain which is present in infancy (Dehaene-Lambertz, Dehaene, & Hertz-Pannier, 2002). In our data, the 5-9 year olds do now show lateralization of information for characters, words, or numbers in VTC (Figure 2), and lateralization increases with age. This suggests the possibility that left-lateralized top-down influences of language areas (Behrmann & Plaut, 2015; Dehaene-Lambertz et al., 2002), as well as white matter changes across development (Takeuchi et al., 2016; Yeatman et al., 2012), generate lateralized distributed representations within VTC in adulthood.

Another novel aspect of our study is its data-driven, computational approach, which enabled us to quantify which voxels are informative and contribute to behavior. This approach revealed that (i) development of a subset of discriminative voxels within lateral VTC is what increases both word and character information from age 5 to adulthood, and (ii) discriminative voxels contain more information than the entire lateral VTC (Figure 4). These findings show for the first time that development of literacy affects a larger set of neuronal populations within VTC, not only just the VWFA (Ben-Shachar et al., 2011; Cantlon et al., 2011; Dehaene et al., 2010). This finding has important implications for assessment of the neural bases of reading disabilities as it suggests that future investigations should consider examining functional differences and atypical development across VTC beyond the VWFA.

Crucially, our results show that the amount of character and word information in lateral VTC predicts reading performance. First, within lateral VTC, discriminative voxels, but not the rest of lateral VTC, predict reading ability. Second, within the set of discriminative voxels, information distributed across voxels with positive preference (i.e., selective voxels) rather than negative preference predicts reading ability.

These findings have two important implications. First, our data advance understanding of the neural basis of reading ability by providing for the first time a neural mechanism explaining why reading ability improves. Notably we found that neural development is driven both by increased within-domain similarity of distributed responses as well as increased between-domain distinctiveness (Figure 3). Together these developments lead to increased information about words in the left lateral VTC, which, in turn, improves reading ability.

Second, our innovative approach provides a new computational and data-driven method to assess which voxels (features) within distributed patterns contribute to behavior. By combing brain and behavioral measurements we show that distributed information across the subset of word/character selective voxels predicts reading ability even as distributed responses across a broader set of discriminative lateral VTC voxels shows substantial neural development. This observation highlights that information in distributed neural responses does not guarantee that it is behaviorally relevant. Importantly, our new approach can be applied broadly across the brain to evaluate the contribution of distributed responses to behavior, and resolves outstanding debates about the utility of MVPA for fMRI data (Dubois, de Berker, & Tsao, 2015; Norman, Polyn, Detre, & Haxby, 2006). Together, our data underscores that it is necessary to combine brain and behavioral measurements to understand the impact of distributed information on behavior, and provides empirical support for the domain-specific view that the left VWFA is critical for reading (Gaillard et al., 2006).

Our data also generate new questions for future research. *First, why does word information develop more than number information in lateral VTC even as both types of stimuli are learned during school years?* It is possible that differences in holistic processing of words vs numbers underlie differences in development of distributed responses to these stimuli. For example, as an adult it is difficult for you not to read the entire pseudoword ‘*Sib*’ in Figure 1a; in contrast, you have likely processed separately each digit of ‘*1453*’ in Figure 1a to infer its quantity. Holistic processing of words may be accomplished via spatial integration by large and foveal population receptive fields (pRFs), in word-selective voxels in left lateral VTC, which continue to develop after age 5 (Gomez et al., 2018). However, if processing of numbers does not involve the same spatial integration as words, number pRFs, and in turn, distributed representations to numbers may show lesser development.

Second, *what features of word/character information develop?* Does learning to read affect the tuning to characters, orthographic information, lexical information, and/or the tuning to the statistics of the language (e.g., bigram frequency, Binder et al., 2006; Vinckier et al., 2007; Glezer, Jiang and Riesenhuber, 2009; Taylor, Rastle and Davis, 2014)? This question can be addressed in the future using fMRI-adaptation (Grill-Spector et al., 1999; Natu et al., 2016; Nordt, Hoehl, & Weigelt, 2016).

Third, *how does the development of word/character information affect development of information to other categories?* Our data show that as domain information for characters develops in VTC, there is no development of information for the domains of bodies, objects, and places. However, consistent with prior research (Golarai et al., 2017) we found developmental increases in domain information to faces. One possibility is that development of face and word information occur in tandem, but are independent from each other. Another possibility, suggested by developmental theories (Behrmann & Plaut, 2015; Dehaene, Cohen, Morais, & Kolinsky, 2015) is that competition on foveal resources for faces and words, produces an interactive development across domains, resulting in the left lateralization of word information and the right lateralization of face information (Behrmann & Plaut, 2015; Dehaene et al., 2015). Future longitudinal research can examine if and how this competition may shape VTC responses.

Finally, *is there a critical period in which development of distributed patterns in VTC can occur, or is this flexibility maintained throughout the lifespan?* Studies of people who gained literacy in adulthood show lesser changes in VWFA response amplitudes with literacy as compared with people who gained literacy as children (Dehaene et al., 2010, 2015). Additionally, research in non-human primates reveal that extensive symbol training leads to development of regions selective to trained symbols in juvenile, but not adult macaque monkeys (Srihasam, Mandeville, Morocz, Sullivan, & Livingstone, 2012). These data suggest that distributed representations to characters and symbols may be more malleable in childhood than adulthood. This hypothesis can be examined in future longitudinal investigations.

In sum, our data show that not only the amount of word information in VTC increases as children learn to read in an anatomical- and hemisphere-specific way, but also that this development is correlated with reading ability. These findings suggest that development of distributed responses to words and characters may be influenced both by foveal biases and interactions with left-lateralized language areas, consequently, they have important implications for both developmental theories and for understanding reading disabilities.

## Methods

### Participants

66 participants including 20 children (ages 5-9 years), 15 preteens (ages 10-12 years), and 31 adults (ages, 22-28 years) participated in this study. Data of 5 children, 1 preteen, and 5 adults were excluded because participants had motion values larger than 2.1 voxels in one or more of the three runs; data of 2 children were excluded because they completed less than 3 fMRI runs; and data from 1 child and 1 preteen were excluded because they fell asleep during fMRI. In total, we report data from 51 participants including 12 children (ages 5-9 years; 11 female), 13 preteens (ages 10-12 years; 6 female), and 26 adults (ages, 22-28 years; 10 female). Participants had normal or corrected-to normal vision and were healthy and typical. The study protocol was approved by the Stanford Internal Review Board on Human Subjects Research. Adult participants and parents of the participating children gave written consent to study participation and children gave written assent.

Participants underwent anatomical and functional MRIs. Prior to MRI, children were trained in a scanner stimulator. Children were invited to MRI sessions only after having successfully completed the scanner simulator training. After completion of MRIs, subjects participated in behavioral tests outside the scanner. Different measurements were performed on different days.

### MRI data acquisition

MRI data were collected at the Stanford Center for Cognitive and Neurobiological Imaging using a 3T Signa scanner (GE Healthcare) and a custom-built phase-array 32-channel receive-only head coil.

#### Anatomical MRI

Whole-brain, high-resolution anatomical scans were acquired using T1-weighted quantitative MRI (qMRI, Mezer et al., 2013), using a spoiled gradient echo sequence with multiple flip angles (α = 4, 10, 20, and 30°; TR=14 ms; TE=2.4 ms. Voxel size= 0.8mm × 0.8mm × 1mm, resampled to 1mm^3^ isotropic. Additionally, we acquired T1-calibration scans using spin-echo inversion recovery with an echo-planar imaging read-out, spectral spatial fat suppression, and a slab inversion pulse (TR=3 s, echo time=minimum full, 2×acceleration, inplane resolution=2 mm^2^; slice thickness= 4 mm).

#### Functional MRI

fMRI data were obtained with a multi-slice EPI sequence (multiplexing factor=3; 48 slices oriented parallel to the parieto-occipital sulcus; TR=1s; TE=30ms; flip angle=76°; FOV = 192 mm; 2.4mm isotropic voxels; one-shot T2*-sensitive gradient echo sequence).

#### fMRI 5 domain/10 category localizer

Participants completed 3 runs of the fMRI experiment. Each run lasted 5 min and 24 s. During fMRI participants viewed images from five domains, each consisting of two categories: characters (pseudowords and numbers), faces (adult and child faces), bodies (headless bodies and limbs), objects (cars and guitars), and places (houses and corridors) as in our prior studies (Stigliani, Weiner, & Grill-Spector, 2015). Pseudowords are the same as in (Glezer et al., 2009; Glezer, Kim, Rule, Jiang, & Riesenhuber, 2015) and have similar bi-gram and trigram frequency as typical English words. Images were grayscale and contained a phase-scrambled background generated from randomly selected images (Figure 1a). Images were presented at a rate of 2Hz, in 4s trials, and did not repeat. Image trials were intermixed with gray luminance screen baseline trials. Trials were counterbalanced across categories and baseline.

<u>*Task:*</u> Participants were instructed to view the images as they fixated on a central dot, and press a button when an image with only the phase-scrambled background appeared. These images appeared randomly 0, 1, or 2 times within a trial.

### Data analysis

Data were analyzed using MATLAB 2012b and mrVista (http://github.com/vistalab) as in previous publications(Gomez, Barnett, et al., 2017; Natu et al., 2016; Stigliani et al., 2015).

#### Anatomical data analysis

An artificial T1-weighted anatomy was generated from qMRI data using mrQ (https://github.com/mezera/mrQ). Brain anatomy was segmented into gray-white matter with FreeSurfer 5.3 (https://surfer.nmr.mgh.harvard.edu/), and manually-corrected to generate cortical surface reconstructions of each participant.

#### Anatomical definition of lateral and medial ventral temporal cortex

Lateral and medial ventral temporal cortex (VTC) were individually defined on each participant’s inflated cortical surface in each hemisphere as in previous publications (Weiner & Grill-Spector, 2010) (Figure 1b). VTC definition: *anterior border*: anterior tip of the mid fusiform sulcus (MFS) which aligned with the posterior end of the hippocampus; *posterior border:* posterior transverse collateral sulcus (ptCoS); *lateral VTC:* extended from the inferior temporal gyrus (ITG) to the MFS; *medial VTC:* extended from the MFS to the medial border of the collateral sulcus (CoS). VTC ROIs were drawn by BJ, divided into the lateral and medial part by MN, and checked by KGS.

#### fMRI data analysis

Data were processed aligned to each participant’s native brain anatomy and were not spatially smoothed. We motion-corrected data within a run and then across runs. Subjects with 3 runs with motion <2.1 voxels were included in the analysis. After exclusion of 8 children, 2 preteens, and 5 adults, there were no significant differences in motion either within or between runs across age groups (Figure 1c). The time courses of each voxel were transformed into percentage signal change and a general linear model (GLM) was fit to each voxel to estimate the contribution of each of the 10 conditions.

#### Multivoxel Pattern Analysis

In each anatomical ROI, multivoxel patterns (MVPs) for each category were represented as a vector of response amplitudes estimated from the GLM in each voxel. These values were transformed to z-scores by subtracting in each voxel its mean voxel response and dividing 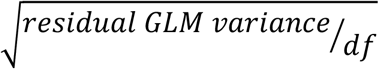 (df=degrees of freedom). To evaluate similarity between MVPs we calculated all pair-wise correlations between MVP pairs resulting in a 10×10 cross-covariance matrix, referred to as representational similarity matrix (RSM, Kriegeskorte, 2008). Each cell in the RSM reflects the average correlation among 3 permutations of MVP pairs (run 1&2, run 2&3, and run 1&3).

#### Winner-take-all classifier

To evaluate category information in MVPs we used an independent winner-take-all (WTA) classifier. We implemented two versions of the WTA classifier. The first, evaluated domain information (characters/faces/bodies/objects/places, chance level 20%). The second, evaluated category information (pseudowords, numbers, adult faces, child faces, headless bodies, limbs, cars, guitars, corridors, buildings, change level 10%). The classifier was trained in each subject and ROI with data from one run and tested how well it predicted the stimulus of interest the subject viewed from MVPs from each of the other two runs. This resulted in six training and testing combinations per condition. We averaged across these combinations for each subject, and then averaged across subjects of each age group.

#### Information in selective and non-selective voxels in lateral VTC

We compared classification performance in selective and non-selective voxels using two analyses (1) lateral VTC character-selective (words+numbers> faces, bodies, objects, places, t>3, voxel level) vs. the remaining voxels, which we refer to as non-character-selective voxels, and (2) lateral VTC word-selective (words > numbers, faces, bodies, objects, places, t>3, voxel level) vs. the remaining voxels which we refer to as non-word-selective voxels.

#### Information in systematically increasing proportions of lateral VTC voxels

We tested how the number of voxels within lateral VTC voxels affects classification in two analyses using different voxel sortings. Analyses were performed for both word classification and character classification as described above. Sorting 1: voxels were sorted by *selectivity*– from highest to lowest *t-value* for the relevant contrast. Sorting 2: voxels were sorted *by distinctiveness*– from highest to lowest *absolute t-value* for the relevant contrast. After sorting of voxels, analyses were identical: We calculated classification performance for increasing portions of lateral VTC voxels according to each sorting, starting with 10% of voxels, increasing by increments of 10%, up to all voxels.

To compare classification across age groups, we fitted each subject’s classification performance as a function of number of voxels a quadratic function, then determined its maximum. Statistical analyses determined if the maximal classification significantly varied across age groups using rmANOVAs on the estimated classification maxima. Due to numerical fitting, the estimated maximum could exceed 100% even as classification performance cannot exceed 100%. Thus, we performed a control analysis in which we rectified the maximum value to 100%. Results did not significantly differ from the original analysis.

### Assessing reading ability

A subset of 7 children, 11 preteens, and 19 adults also completed the *word identification* and *word attack* tests from the Woodcock Reading Mastery Test (WRMT) outside the scanner. In the *word identification task*, participants were instructed to read a list of words as accurately as possible. In the *word attack task*, were instructed to read a list of pseudowords as accurately as possible. Tests do not have a time limit, but end when the participant makes four consecutive errors or has completed reading the list.

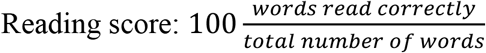

#### Relating reading ability to word and character information in VTC

We measured if there was a significant correlation between participant’s reading ability as measured by the WRMT and information in lateral VTC. Correlations were measured between each reading test score (word identification/word attack test) and each classification (character/word) from lateral VTC. Significant correlations were followed with a subsequent analysis in which age was included as a factor. We report if correlations remained significant if age is partialled out of the correlation.

Since analyses of information in lateral VTC revealed that a subset of voxels in VTC contribute to either character or word information, we performed correlation analyses between reading and subsets of lateral VTC voxels: (1) entire lateral VTC, (2) 30% of most discriminative lateral VTC voxels, which yielded best word classification (Figure 4), and (3) the remaining nonword-distinctive lateral VTC voxels (Figure 4). The same analyses were done for 30% of lateral VTC voxels that where most discriminative for characters (Figure 4, Figure S4). We evaluated if the correlations in (2) and (3) were significantly different and within (2) if information in most or least selective voxels correlated with reading ability.

### Statistical analyses

Unless otherwise noted, statistical analyses included the whole sample (5-9, n = 12; 10-12, n = 13; 22-28, n = 26). Statistical analyses were done using MATLAB 2015a. Outliers in boxplots are defined as values that were more than 1.5 times away from the interquartile range from the top or bottom of the box and are indicated by black dots.

In analyses related to Figure 1c we tested if age groups differed significantly in the amount of motion during scanning using a repeated measures analysis of variance (rmANOVA) with factors of age group (5-9/10-12/22-28) and motion type (within-run motion, between-run motion). Similarly, in analyses related to Figure 1d we tested in each partition (lateral VTC, medial VTC) if there were statistically significant between group differences in the number of voxels using ANOVAs with the factor age group (5-9/10-12/22-28). The same procedure was applied for analyses related to Figure 1e, in which we tested the statistical significance of the number of word-selective voxels across groups.

In analyses related to Figure 2 we first tested if classification performance for characters was significantly different from chance level (20%) in each age group using one-sample t-tests. Next, we tested for significant differences in classification performance using a 3-way rmANOVA with factors of age group (5-9/10-12/22-28), partition (lateral VTC/medial VTC), and hemisphere (left/right). Similar analyses were run for classification performance for character types, i.e. for pseudoword and number classification. Here, classification performance was tested against the chance level of 10%. Significant differences in decoding performance were tested using a 4-way rmANOVA with factors of age group (5-9/10-12/22-28), character type (numbers/pseudowords), partition (lateral VTC/medial VTC), and hemisphere (left/right). To follow up on significant interactions between character type and age group we conducted separate 3-way rmANOVAs on number and pseudoword classification with factors of age group (5-9/10-12/22-28), partition (lateral VTC/medial VTC) and hemisphere (left/right). To further follow up on the significant effect of age, we directly compared classification performance for words across age groups using post-hoc t-tests.

In analyses related to Figure 3 we first tested if the mean within-domain correlations for pseudowords and numbers across partitions and hemispheres were significantly different from zero using one-sample t-tests. Next, we tested for significant differences of correlations using 3 rmANOVAs. The rmANOVAs tested correlations (i) within the domain of characters (w-w; b-n) with factors of age group (5-9/10-12/22-28), partition (lateral VTC/medial VTC), hemisphere (left/right), and character type (w-w/n-n), (ii) across character types (w-n), with factors of age group (5-9/10-12/22-28), partition (lateral VTC/medial VTC), and hemisphere (left/right) and (iii) between-domains (w-nw; n-nn) including the factors of age group (5-9/10- 12/22-28), partition (lateral VTC/medial VTC), hemisphere (left/right), and character type (w-nw/n-nn). Furthermore, we tested for each character type in each partition and hemisphere if there was a significant effect of age group (1-way ANOVAs with the factor of age group (5- 9/10-12/22-28)) to follow up on significant interactions revealed in rmANOVAs measured above. Significant effects of age are shown in Figure 3 with asterisks.

In analyses related to Figure 4a we tested differences in word classification in lateral VTC in word-selective and non-selective voxels using a 3-way rmANOVA with factors of age group (5-9/10-12/22-28), hemisphere (left/right), and voxel type (selective/non-selective).

In analyses related to Figure 4b we compared the estimated maximal classification performance by populations of voxels sorted by selectivity using a 2-way rmANOVA with factors of age group (5-9/10-12/22-28) and hemisphere (left/right). Same analyses were conducted on classification performance on voxels sorted voxels by distinctiveness for the data shown in Figure 4c. Corresponding analyses were performed for character classification in character-selective and non-selective voxels in Figure 4d-f. We further tested if word information developed also in the remainder of non-selective/non discriminative voxels. Thus, we ran an additional rmANOVA with factors of age group (5-9/10-12/22-28) and hemisphere (left/right) on word decoding in the remaining voxels (i.e. the lateral VTC voxels except the set of voxels which generated the maximal classification). In four subjects, classification performance was identical for all voxel set sizes exceeding 10% of the lateral VTC. For these subjects, we included the minimal set (10% of voxels) as the ones achieving maximal classification, and tested classification in the remaining 90% of voxels.

In analyses related to Figure 5 we tested if reading scores (WRMT) are correlated with classification performance for words and/or characters by calculating the Pearson correlation coefficient between these values and their significance. If correlations were found to be significant, we performed an additional partial correlation analysis to control for the effect of age. We also tested if correlations between reading scores and classification in different subsets of lateral VTC voxels (30% of most discriminative lateral VTC voxels vs the remaining non-discriminative lateral VTC voxels) differed significantly using (http://quantpsy.org/corrtest/corrtest2.htm), which includes converting the correlation coefficients into z-scores using Fisher’s r-to-z transformation.

### Analysis of V1 MVPs

V1 was defined using data from a separate retinotopic mapping experiment. A subset of participants comprising 8 children ages 5-9, 12 children ages 10-12, and 19 adults took part in a retinotopic mapping experiment using black and white checkerboard bars (width = 2° of visual angle, length = 14°) which changed contrast at a rate of 2Hz, that swept across the screen. During retinotopic mapping subjects were instructed to fixate on a central stimulus and perform a color exchange task. Subjects’ fixations were monitored with an eye tracker. Checkerboard bars swept the visual field in 8 different configurations in each run (4 orientations: 0°, 45°, 90°, 135°, each orientation was swept in 2 directions that were orthogonal to the bar). Same as (Dumoulin & Wandell, 2008; Weiner & Grill-Spector, 2011). We used checkerboard stimuli as they are the most ubiquitous stimuli that is used for population receptive field (pRF) mapping, and do not require cognitive processing that may differ across age groups. Subjects participated in 4 such runs, each run lasted 3 minutes and 24 seconds. Retinotopic data were collected on the same scanner as the main experiment and at the same resolution, with a 16-channel surface coil, acceleration factor x 2, and 28 slices. After fitting the pRF model (Dumoulin & Wandell, 2008) with a compressive spatial summation (Kay, Winawer, Mezer, & Wandell, 2013) in each voxel, maps of pRF phase and eccentricity were projected onto an the inflated cortical surface of each subject’s brain. V1 was defined in each hemisphere as the visual field map containing a hemifield representation in which the horizontal meridian occupied the calcarine sulcus, the lower vertical meridian occupied the upper lip of the calcarine and the upper visual meridian occupied the lower lip of the calcarine.

After defining V1 we performed MVPA and classification analyses on V1 of each hemisphere. This analysis tested whether category information in VTC was beyond low-level information that could be extracted from V1 MVPs. Results shown in Figure S1b indicate that the domain information in VTC is significantly higher than V1 (main effect of area, *F(*2,72)=142.1, *p*<0.001; lateral VTC > V1, *t*(38)=16.32,*p*<0.001; medial VTC > V1, *t*(38)=10.13,*p*<0.001).

## Data and software availability

All code relevant to data analysis for the main findings (Figures 1-5) will be available on github.com/VPNL. Any source data relevant to these analyses will also be made available upon request. The majority of the code used in this study was derived from scripts and functions available through the open-source vistasoft code library: https://github.com/vistalab/vistasoft

## Author contributions

M.N. developed the analysis pipeline, analyzed the data, and wrote the manuscript.

V.N. and J.G. contributed to experimental design, collection and analysis of data, and preparation of the manuscript.

B.J., M.B contributed to collection of data, and preparation of the manuscript.

K.G.-S. designed the experimental design, developed the analysis pipeline, contributed to the data analysis, and wrote the manuscript.

## Acknowledgements

This work was supported by a scholarship of the German National Academic Foundation and by the Ruhr University Research School PLUS, funded by Germany’s Excellence Initiative [DFG GSC 98/3] awarded to MN, NIH grants 1RO1EY02231801A1, 1RO1EY02391501A1 to KGS, training grant 5T32EY020485 supporting VN, and the NSF Graduate Research Development Program (grant DGE-114747) and Ruth L. Kirschstein National Research Service Award (grant F31EY027201) to JG.

## Supplemental Figures

**Figure S1.**
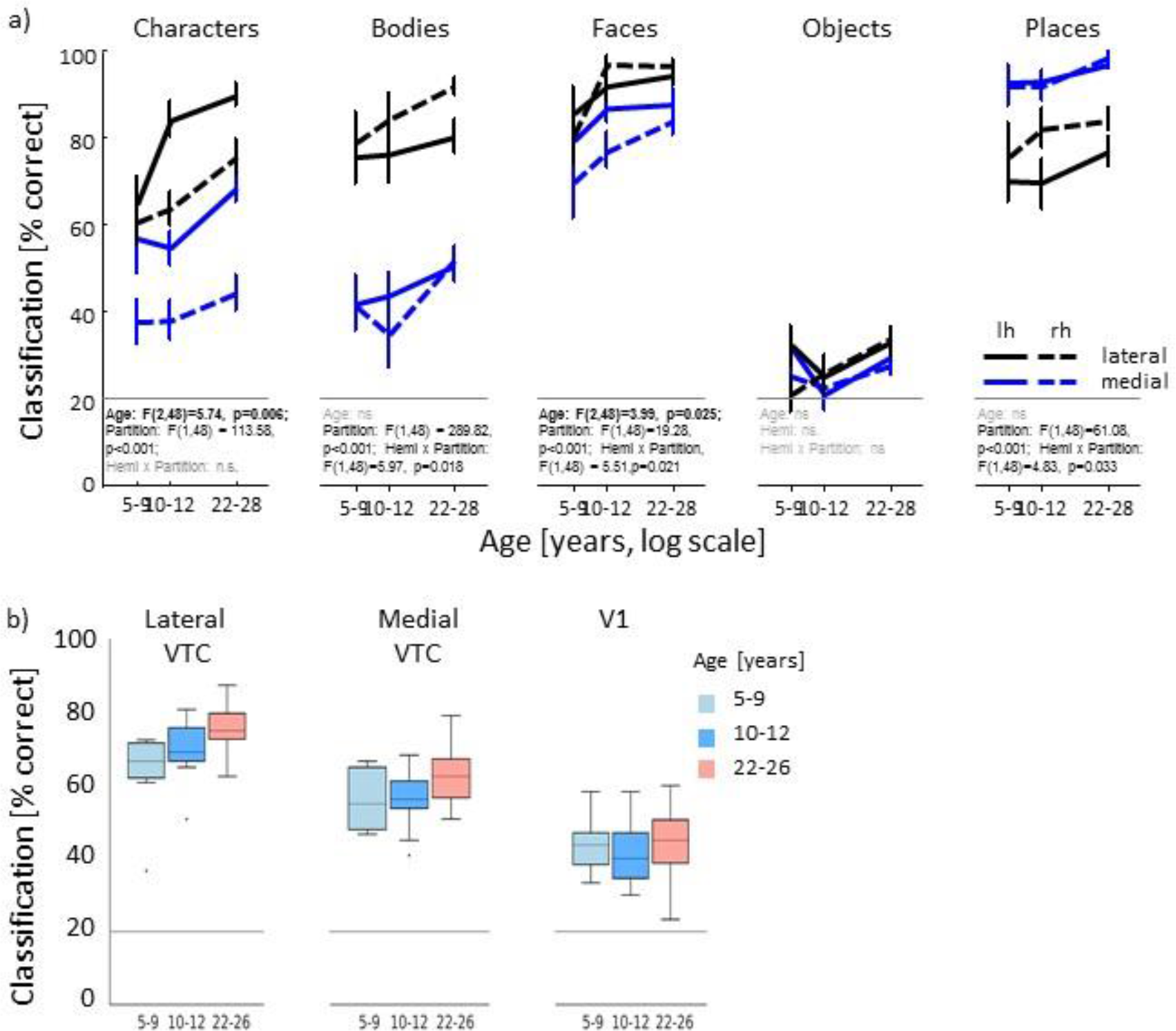
Related to Figure 2. Differential development of domain information in VTC and classification performance in VTC partitions and V1. (a) Decoding performance of domains (characters, bodies, faces, objects, and places) from VTC multivoxel patterns (MVPs) using a winner-take-all (WTA) classifier, *y-axis:* classification performance; x-axis: age groups (5-9 year olds, *n* = 12; 10-12 year olds, *n* = 13; 22-28 year olds, *n* = 26) ticks are at the mean age of each group displayed on a logarithmic scale. *Block:* lateral VTC data. *Blue:* medial VTC data. *Solid:* left hemisphere. *Dashed:* right hemisphere. *Horizontal gray line:* Chance level (20%). *Error bars:* standard error of the mean (SEM). We tested for developmental differences in classification performance using 3-way rmANOVAs with factors age group (5-9/10- 12/22-28), partition (lateral VTC/medial VTC), and hemisphere (left/right) in classification of each of the 5 domains. We report for all domains age, partition, and partition by hemisphere interactions as these were significant for some domains, (b) Mean domain classification performance in each age group in lateral VTC, medial VTC, and VI. Classification performance of the WTA is shown averaged across hemispheres and domains. Data are from a subset of participants, who participated in a retinotopy experiment. *Light blue:* children (5-9 year olds, n =8); *Blue:* preteen (10-12 year olds, *n* = 12); *Orange:* adults (22-28 year olds, *n =*19). *Horizontal gray line:* Chance level (20%). Horizontal lines in the boxplots indicate the median. The whiskers correspond to the range of the data excluding outliers, which include data within ± 2.7 standard deviations from the mean and include 99.3% of the data. Statistical comparison of domain classification in the three ROIs used a 2-way rmANOVA on classification performance with factors of age group (5-9/10-12/22-28) and ROI (lateral VTC. medial VTC, VI). Classification of domain information was not due to low-level features, as decoding domain information from VTC was significantly higher than from VI (main effect of ROI, *F*2,72)=142.1, p<0.001). Additionally, development occurred in VTC but not VI (ROI x age interaction, *F*(4,72) = 2.78, *p* = 033).

**Figure S2.**
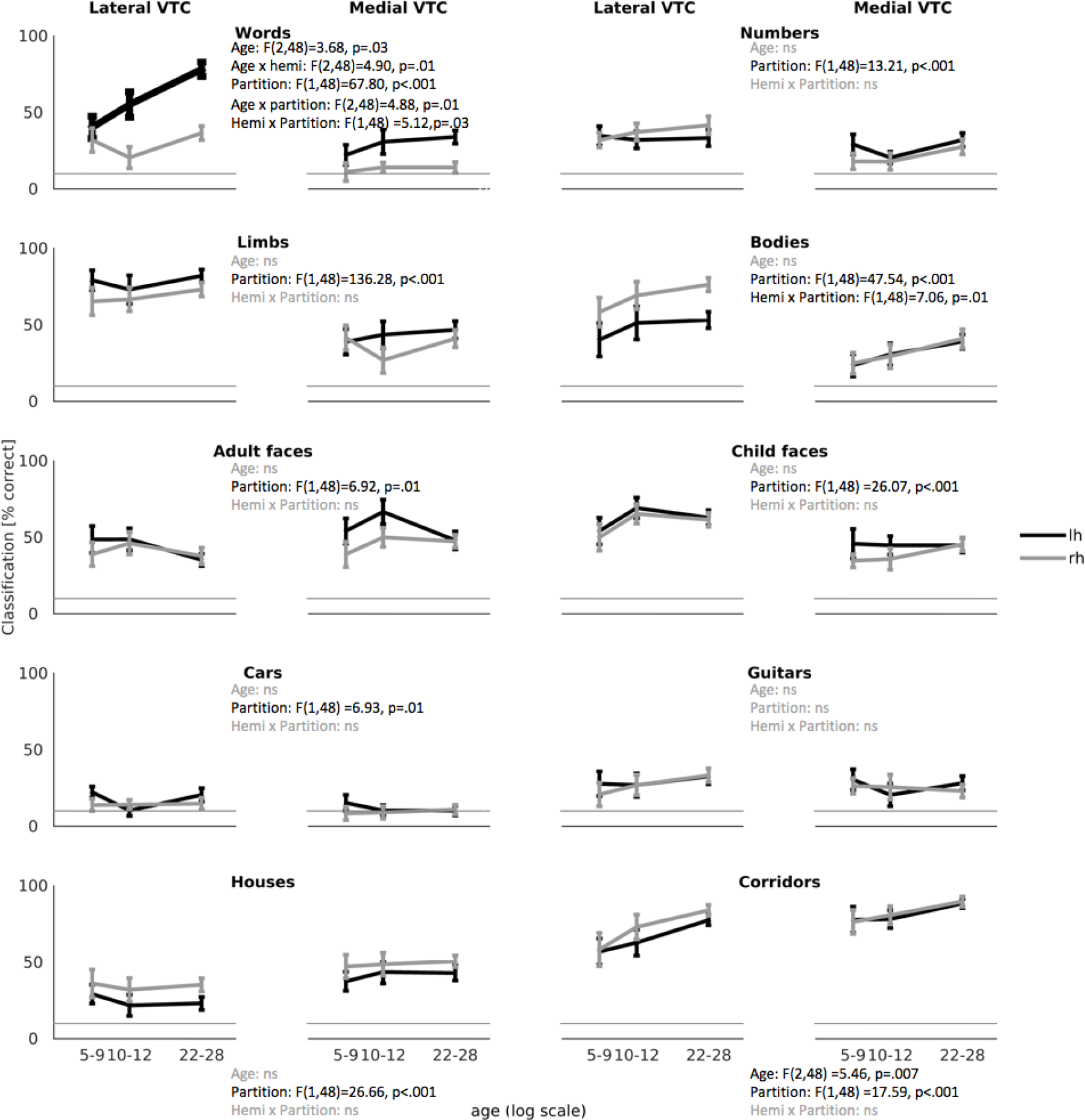
Related to Figure 2. Differential development of category type information in VTC. Classification performance of each category type from VTC multivoxel patterns (MVPs) using a WTA classifier. Each row presents data from one domain, and each category is listed above each plot, *y-axis:* classification performance; x-axis: age groups (5-9 year olds, *n* = 12; 10-12 year olds, *n* = 13; 22-28 year olds, *n* = 26) ticks are at the mean age of each group displayed on a logarithmic scale. *Black:* left hemisphere. *Gray:* right hemisphere. *Horizontal gray line:* Chance level (10%). *Error bars:* standard error of the mean. Results of a 3-way ANOVA with factors of age-group (5-9/10-12/22-28), partition (lateral VTC/medial VTC), and hemisphere (left/right) on classification performance for each category are displayed for each category. We report for all categories age, partition, and partition by hemisphere interactions as these were significant for some categories. All other interactions were not significant.

**Figure S3.**
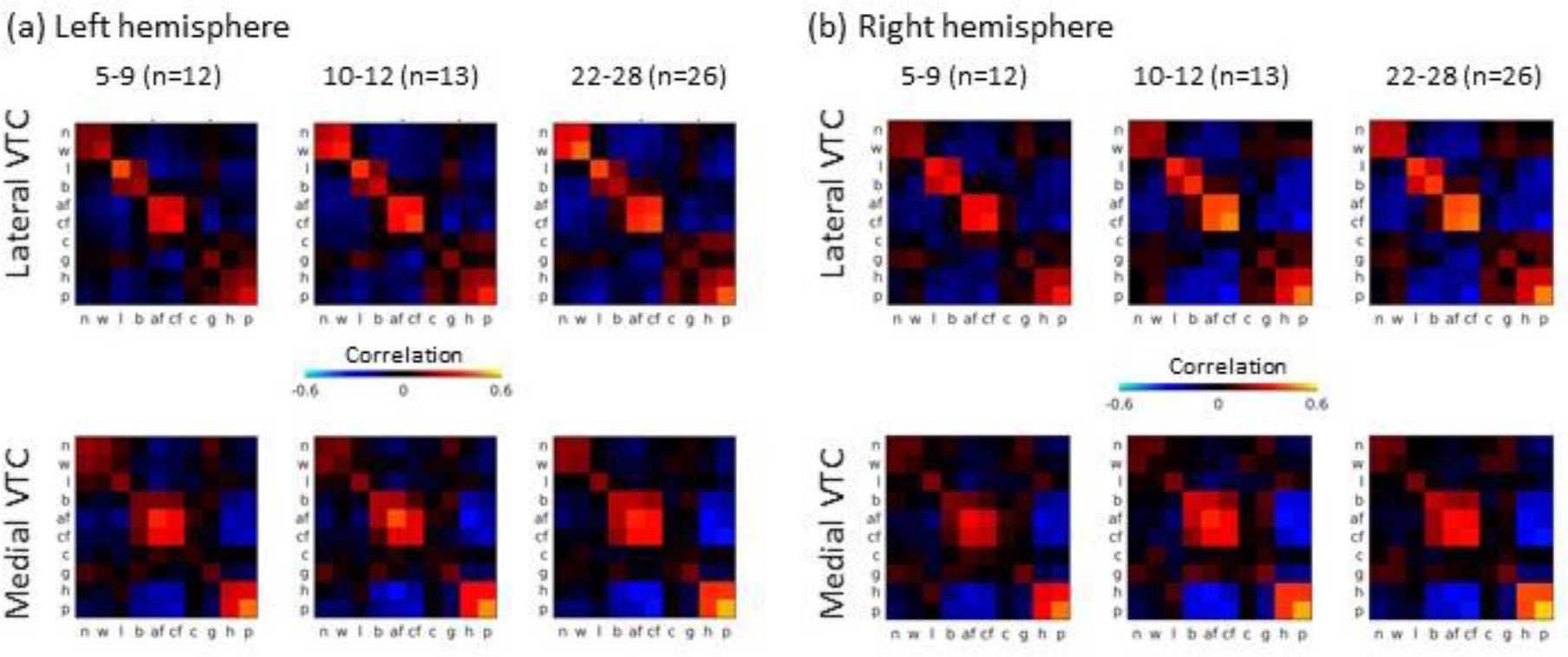
Related to Figure 3. Representational similarities of visual categories in VTC. Representational similarity matrices (RSM) of distributed responses across (a) left and (b) right VTC in children and adults. The matrices display the Pearson correlation coefficients between multivoxel patterns for 10 stimulus categories from all unique combination of runs, averaged across combinations of runs and subjects. The RSM on the left display data from children (5-9 year olds, *n =*12), the RSM in the middle display data from preteens (10-12 year olds, *n =*13) and the RSM on the right show data from adults (22-28 year olds, *n =*26). The top row shows data from lateral VTC and the bottom row depicts data from medial VTC. Acronyms for 10 categories are indicated as follows: n = numbers, w = words (pseudowords), I = limbs, b = bodies, af = adult faces, cf = child faces, c = cars, g = guitars, h = houses, p = places (corridors).

**Figure S4.**
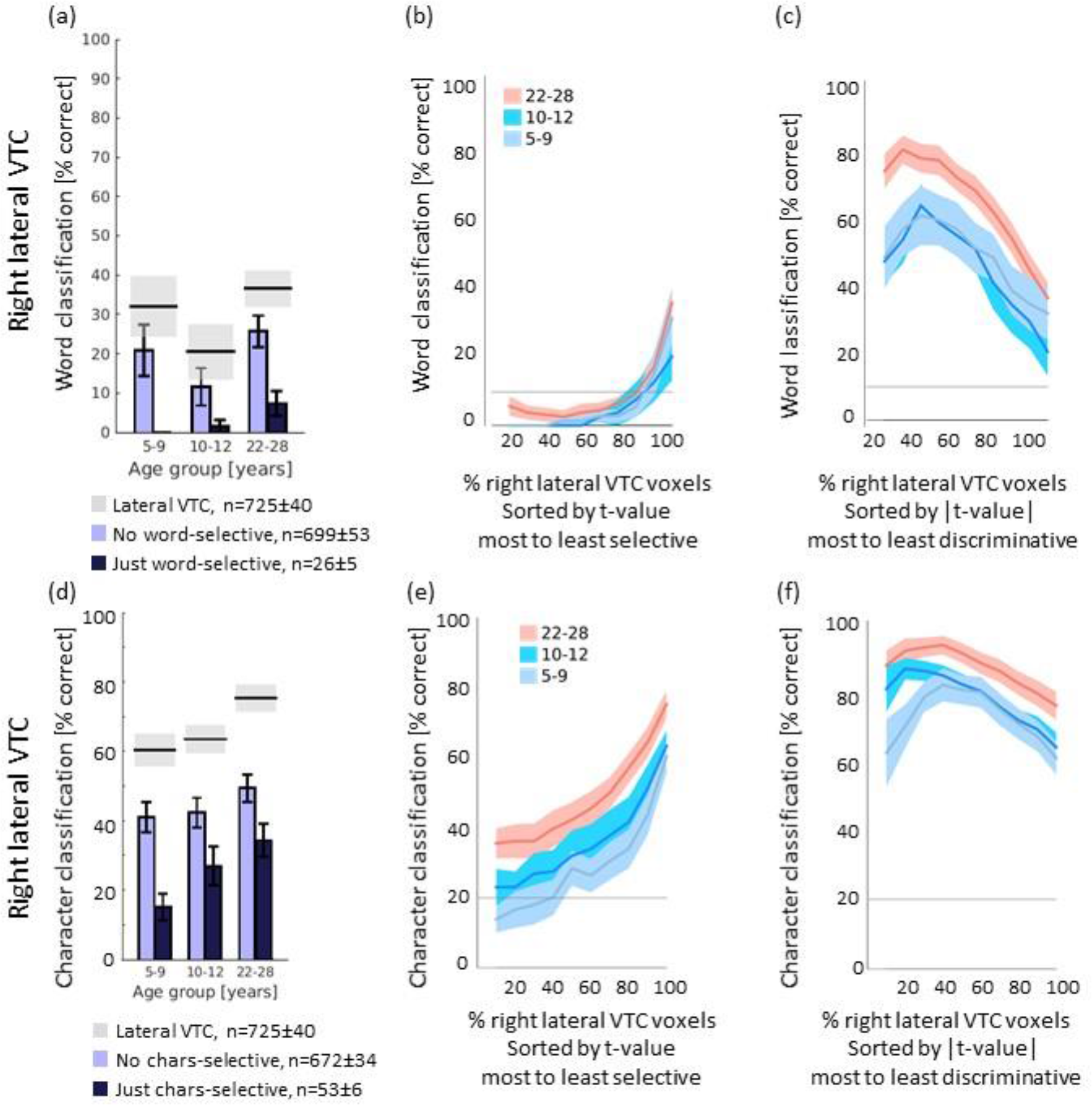
Related to Figure 4. Development of word and character Information in right VTC. (a) *Dark blue:* Mean classification performance of word-selective voxels within right lateral VTC (t>3). *Light blue:* Mean classification performance of right lateral VTC voxels excluding the word-selective voxels (t < 3); *Horizontal lines:* mean (black) and SEM (gray) classification performance using the entire right lateral VTC. *Legend:* number of voxels included in each analysis. As not all subjects had word-selective voxels, only the subjects that had word-selective voxels are included (5-9 year olds, *n* =9; 10-12 year olds, *n* = 11; 22-28 year olds, n = 25). (b) Word classification performance as a function of the percentage of right lateral VTC voxels sorted by their preference to words (from most to least selective, that is descending t-value for the contrast words>non-words). *Lines:* mean performance; *Shaded areas:* SEM. (c) Word classification performance as a function of percentage of right lateral VTC voxels sorted by their discriminative value (from most to least discriminative, that is descending |t-value| for the contrast words>non-words). (d,e,f) Same as a-c but for character information, (d) As not all subjects had character-selective voxels, only the subjects that had character-selective voxels are included (5-9 year olds, *n =*11; 10-12 year olds, *n* = 13; 22-28 year olds,n= 26).

**Figure S5.**
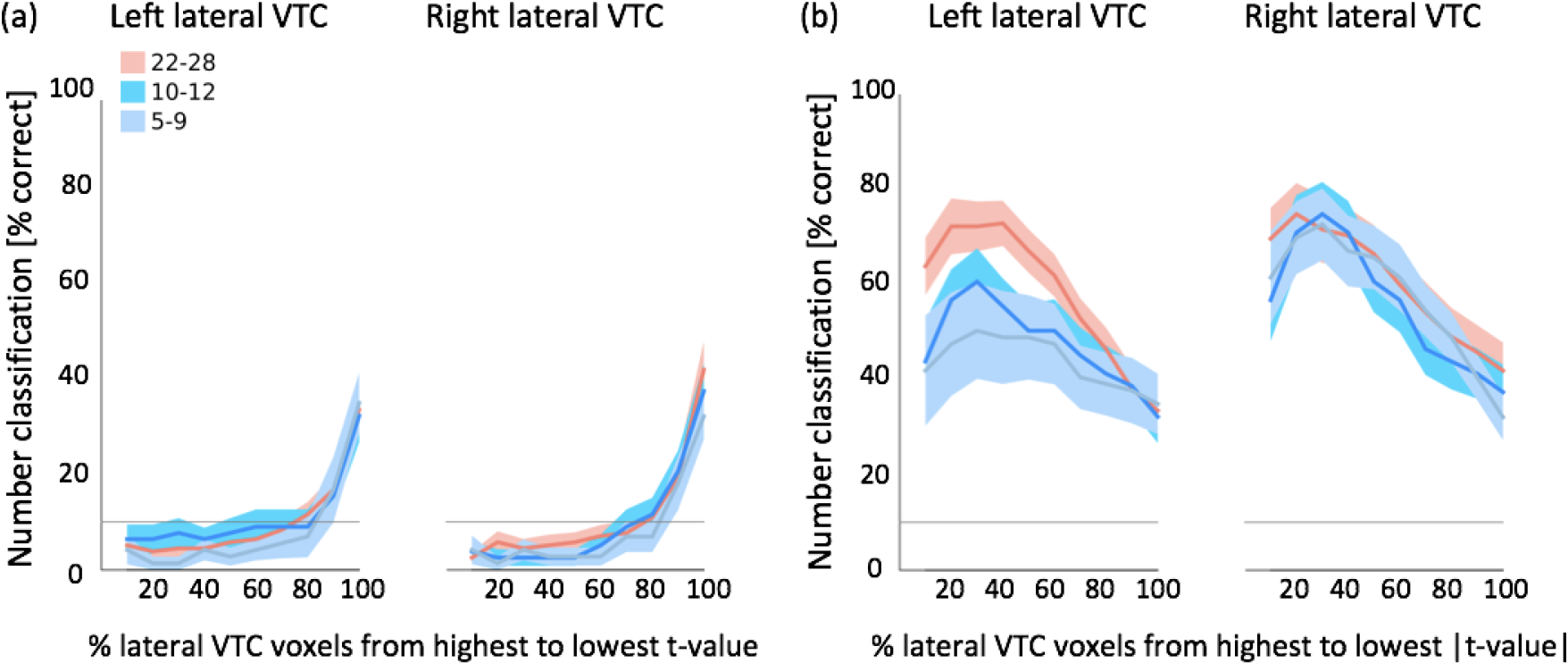
Related to Figure 4. Number classification in selective and discriminative voxels. (a, b) Mean classification performance of number information as function of proportion VTC voxels in each age group in left and right lateral VTC. *Shaded areas:* SEM. (a) Voxels are sorted from most to least number-selective (descending t-value for the contrast numbers>non-numbers). (b) Voxels are sorted from most to least number discriminative (descending | t-value | for the contrast numbers>non-numbers).

## References

Behrmann, M., & Plaut, D. C. (2015). A vision of graded hemispheric specialization. Annals of the New York Academy of Sciences, 1359(1), 30–46. https://doi.org/10.1111/nyas.12833

Ben-Shachar, M., Dougherty, R. F., Deutsch, G. K., & Wandell, B. A. (2011). The development of cortical sensitivity to visual word forms. Journal of Cognitive Neuroscience, 23(9), 2387–2399.

Ben-Shachar, M., Dougherty, R. F., & Wandell, B. A. (2007). White matter pathways in reading. Current Opinion in Neurobiology, 17(2), 258–270. https://doi.org/10.1016/j.conb.2007.03.006

Binder, J. R., Medler, D. A., Westbury, C. F., Liebenthal, E., & Buchanan, L. (2006). Tuning of the human left fusiform gyrus to sublexical orthographic structure. Neuroimage, 33(2), 739–748.

Cantlon, J. F., Pinel, P., Dehaene, S., & Pelphrey, K. A. (2011). Cortical representations of symbols, objects, and faces are pruned back during early childhood. Cerebral Cortex, 21(1), 191–199. https://doi.org/10.1093/cercor/bhq078

Carlson, T., Tovar, D. A., Alink, A., & Kriegeskorte, N. (2013). Representational dynamics of object vision: The first 1000 ms. Journal of Vision, 13(10), 1. Retrieved from http://dx.doi.org/10.1167/13.10.1

Carreiras, M., Seghier, M. L., Baquero, S., Estévez, A., Lozano, A., Devlin, J. T., & Price, C. J. (2009). An anatomical signature for literacy. Nature, 461(7266), 983–986. https://doi.org/10.1038/nature08461

Cohen, L., Dehaene, S., Naccache, L., Lehéricy, S., Dehaene-Lambertz, G., Hénaff, M.-A., & Michel, F. (2000). The visual word form areaSpatial and temporal characterization of an initial stage of reading in normal subjects and posterior split-brain patients. Brain, 123(2), 291–307. https://doi.org/10.1093/brain/123.2.291

Connolly, A. C., Guntupalli, J. S., Gors, J., Hanke, M., Halchenko, Y. O., Wu, Y. C., Abdi, H., & Haxby, J. V. (2012). The representation of biological classes in the human brain. The Journal of Neuroscience: The Official Journal of the Society for Neuroscience, 32(8), 2608–2618. https://doi.org/10.1523/JNEUROSCI.5547-11.2012

Cox, D. D., & Savoy, R. L. (2003). Functional magnetic resonance imaging (fMRI) “brain reading”: detecting and classifying distributed patterns of fMRI activity in human visual cortex. NeuroImage, 19(2), 261–270. https://doi.org/10.1016/S1053-8119(03)00049-1

Dehaene-Lambertz, G., Dehaene, S., & Hertz-Pannier, L. (2002). Functional neuroimaging of speech perception in infants. Science, 298(5600), 2013–2015.

Dehaene, S., Cohen, L., Morais, J., & Kolinsky, R. (2015). Illiterate to literate: behavioural and cerebral changes induced by reading acquisition. Nature Reviews Neuroscience, 16(4), 234–244.

Dehaene, S., Le Clec’H, G., Poline, J.-B., Le Bihan, D., & Cohen, L. (2002). The visual word form area: a prelexical representation of visual words in the fusiform gyrus. Neuroreport, 13(3), 321–325.

Dehaene, S., Pegado, F., Braga, L. W., Ventura, P., Nunes Filho, G., Jobert, A., Dehaene-Lambertz, G., Kolinsky, R., Morais, J., & Cohen, L. (2010). How learning to read changes the cortical networks for vision and language. Science, 330(6009), 1359–1364.

Dubois, J., de Berker, A. O., & Tsao, D. Y. (2015). Single-Unit Recordings in the Macaque Face Patch System Reveal Limitations of fMRI MVPA. The Journal of Neuroscience, 35(6), 2791 LP-2802. Retrieved from http://www.jneurosci.org/content/35/6/2791.abstract

Dumoulin, S. O., & Wandell, B. A. (2008). Population receptive field estimates in human visual cortex. Neuroimage, 39(2), 647–660. https://doi.org/10.1016/j.neuroimage.2007.09.034

Gaillard, R., Naccache, L., Pinel, P., Clémenceau, S., Volle, E., Hasboun, D., Dupont, S., Baulac, M., Dehaene, S., Adam, C., & Cohen, L. (2006). Direct Intracranial, fMRI, and Lesion Evidence for the Causal Role of Left Inferotemporal Cortex in Reading. Neuron, 50(2), 191–204. https://doi.org/10.1016/j.neuron.2006.03.031

Glezer, L. S., Jiang, X., & Riesenhuber, M. (2009). Evidence for Highly Selective Neuronal Tuning to Whole Words in the “Visual Word Form Area.” Neuron, 62(2), 199–204. https://doi.org/10.1016/j.neuron.2009.03.017

Glezer, L. S., Kim, J., Rule, J., Jiang, X., & Riesenhuber, M. (2015). Adding Words to the Brain’s Visual Dictionary: Novel Word Learning Selectively Sharpens Orthographic Representations in the VWFA. Journal of Neuroscience, 35(12), 4965–4972. https://doi.org/10.1523/JNEUROSCI.4031-14.2015

Golarai, G., Ghahremani, D. G., Whitfield-Gabrieli, S., Reiss, A., Eberhardt, J. L., Gabrieli, J. D., & Grill-Spector, K. (2007). Differential development of high-level visual cortex correlates with category-specific recognition memory. Nat Neurosci, 10(4), 512–522. Retrieved from http://www.ncbi.nlm.nih.gov/entrez/query.fcgi?cmd=Retrieve&db=PubMed&dopt=Citation&list_uids=17351637

Golarai, G., Liberman, A., & Grill-Spector, K. (2017). Experience Shapes the Development of Neural Substrates of Face Processing in Human Ventral Temporal Cortex. Cerebral Cortex, bhv314. https://doi.org/10.1093/cercor/bhv314

Golarai, G., Liberman, A., Yoon, J. M., & Grill-Spector, K. (2010). Differential development of the ventral visual cortex extends through adolescence. Frontiers in Human Neuroscience, 3, 80. https://doi.org/10.3389/neuro.09.080.2009

Gomez, J., Barnett, M. A., Natu, V., Mezer, A., Palomero-Gallagher, N., Weiner, K. S., Amunts, K., Zilles, K., & Grill-Spector, K. (2017). Microstructural proliferation in human cortex is coupled with the development of face processing. Science, 355(6320), 68–71.

Gomez, J., Natu, V., Jeska, B., Barnett, M., & Grill-Spector, K. (in press). Development differentially sculpts receptive fields across human visual cortex. Nature Communications.

Grill-Spector, K., Kushnir, T., Edelman, S., Avidan, G., Itzchak, Y., & Malach, R. (1999). Differential processing of objects under various viewing conditions in the human lateral occipital complex. Neuron, 24(1), 187–203.

Grill-Spector, K., & Weiner, K. S. (2014). The functional architecture of the ventral temporal cortex and its role in categorization. Nature Reviews. Neuroscience, 15(8), 536–548. https://doi.org/10.1038/nrn3747

Gullick, M. M., & Booth, J. R. (2015). The direct segment of the arcuate fasciculus is predictive of longitudinal reading change. Developmental Cognitive Neuroscience, 13, 68–74. https://doi.org/10.1016/j.dcn.2015.05.002

Hannagan, T., Amedi, A., Cohen, L., Dehaene-Lambertz, G., & Dehaene, S. (2015). Origins of the specialization for letters and numbers in ventral occipitotemporal cortex. Trends in Cognitive Sciences, 19(7), 374–382.

Hasson, U., Levy, I., Behrmann, M., Hendler, T., & Malach, R. (2002). Eccentricity bias as an organizing principle for human high-order object areas. Neuron, 34(3), 479–490. Retrieved from http://www.ncbi.nlm.nih.gov/entrez/query.fcgi?cmd=Retrieve&db=PubMed&dopt=Citation&list_uids=11988177

Haxby, J. V, Gobbini, M. I., Furey, M. L., Ishai, A., Schouten, J. L., & Pietrini, P. (2001). Distributed and overlapping representations of faces and objects in ventral temporal cortex. Science, 293(5539), 2425–2430. Retrieved from http://www.ncbi.nlm.nih.gov/entrez/query.fcgi?cmd=Retrieve&db=PubMed&dopt=Citation&list_uids=11577229

Kay, K. N., Winawer, J., Mezer, A., & Wandell, B. A. (2013). Compressive spatial summation in human visual cortex. Journal of Neurophysiology, 110(2), 481–494. https://doi.org/10.1152/jn.00105.2013

Kriegeskorte, N. (2008). Representational similarity analysis – connecting the branches of systems neuroscience. Frontiers in Systems Neuroscience. https://doi.org/10.3389/neuro.06.004.2008

Levy, I., Hasson, U., Avidan, G., Hendler, T., & Malach, R. (2001). Center-periphery organization of human object areas. Nat Neurosci, 4(5), 533–539. Retrieved from http://www.ncbi.nlm.nih.gov/entrez/query.fcgi?cmd=Retrieve&db=PubMed&dopt=Citation&list_uids=11319563

Malach, R., Levy, I., & Hasson, U. (2002). The topography of high-order human object areas. Trends Cogn Sci, 6(4), 176–184. Retrieved from http://www.ncbi.nlm.nih.gov/entrez/query.fcgi?cmd=Retrieve&db=PubMed&dopt=Citation&list_uids=11912041

Mezer, A., Yeatman, J. D., Stikov, N., Kay, K. N., Cho, N. J., Dougherty, R. F., Perry, M. L., Parvizi, J., Hua le, H., Butts-Pauly, K., & Wandell, B. A. (2013). Quantifying the local tissue volume and composition in individual brains with magnetic resonance imaging. Nature Medicine, 19(12), 1667–1672. https://doi.org/10.1038/nm.3390

Natu, V. S., Barnett, M. A., Hartley, J., Gomez, J., Stigliani, A., & Grill-Spector, K. (2016). Development of Neural Sensitivity to Face Identity Correlates with Perceptual Discriminability. The Journal of Neuroscience: The Official Journal of the Society for Neuroscience, 36(42), 10893–10907. https://doi.org/10.1523/JNEUROSCI.1886-16.2016

Nordt, M., Hoehl, S., & Weigelt, S. (2016). The use of repetition suppression paradigms in developmental cognitive neuroscience. Cortex, 80, 61–75. https://doi.org/10.1016/j.cortex.2016.04.002

Norman, K. A., Polyn, S. M., Detre, G. J., & Haxby, J. V. (2006). Beyond mind-reading: multi-voxel pattern analysis of fMRI data. Trends in Cognitive Sciences, 10(9), 424–430. https://doi.org/https://doi.org/10.1016/j.tics.2006.07.005

Rauschecker, A. M., Bowen, R. F., Perry, L. M., Kevan, A. M., Dougherty, R. F., & Wandell, B. A. (2011). Visual feature-tolerance in the reading network. Neuron, 71(5), 941–953. https://doi.org/10.1016/j.neuron.2011.06.036

Schlaggar, B. L., & McCandliss, B. D. (2007). Development of Neural Systems for Reading. Annual Review of Neuroscience, 30(1), 475–503. https://doi.org/10.1146/annurev.neuro.28.061604.135645

Srihasam, K., Mandeville, J. B., Morocz, I. A., Sullivan, K. J., & Livingstone, M. S. (2012). Behavioral and Anatomical Consequences of Early versus Late Symbol Training in Macaques. Neuron, 73(3), 608–619. https://doi.org/10.1016/j.neuron.2011.12.022

Stigliani, A., Weiner, K. S., & Grill-Spector, K. (2015). Temporal Processing Capacity in High-Level Visual Cortex Is Domain Specific. The Journal of Neuroscience: The Official Journal of the Society for Neuroscience, 35(36), 12412–12424. https://doi.org/10.1523/JNEUROSCI.4822-14.2015

Takeuchi, H., Taki, Y., Hashizume, H., Asano, K., Asano, M., Sassa, Y., Yokota, S., Kotozaki, Y., Nouchi, R., & Kawashima, R. (2016). Impact of reading habit on white matter structure: Cross-sectional and longitudinal analyses. NeuroImage, 133, 378–389. https://doi.org/10.1016/j.neuroimage.2016.03.037

Taylor, J. S. H., Rastle, K., & Davis, M. H. (2014). Distinct Neural Specializations for Learning to Read Words and Name Objects. Journal of Cognitive Neuroscience, 26(9), 2128–2154. https://doi.org/10.1162/jocn_a_00614

Vinckier, F., Dehaene, S., Jobert, A., Dubus, J. P., Sigman, M., & Cohen, L. (2007). Hierarchical Coding of Letter Strings in the Ventral Stream: Dissecting the Inner Organization of the Visual Word-Form System. Neuron, 55(1), 143–156. https://doi.org/10.1016/j.neuron.2007.05.031

Wandell, B. A., Rauschecker, A. M., & Yeatman, J. D. (2012). Learning to see words. Annu Rev Psychol, 63, 31–53. https://doi.org/10.1146/annurev-psych-120710-100434

Weiner, K. S., Golarai, G., Caspers, J., Chuapoco, M. R., Mohlberg, H., Zilles, K., Amunts, K., & Grill-Spector, K. (2014). The mid-fusiform sulcus: A landmark identifying both cytoarchitectonic and functional divisions of human ventral temporal cortex. Neuroimage, 84, 453–465. https://doi.org/10.1016/j.neuroimage.2013.08.068

Weiner, K. S., & Grill-Spector, K. (2010). Sparsely-distributed organization of face and limb activations in human ventral temporal cortex. Neuroimage, 52(4), 1559–1573. Retrieved from http://www.ncbi.nlm.nih.gov/entrez/query.fcgi?cmd=Retrieve&db=PubMed&dopt=Citation&list_uids=20457261

Weiner, K. S., & Grill-Spector, K. (2011). Not one extrastriate body area: Using anatomical landmarks, hMT+, and visual field maps to parcellate limb-selective activations in human lateral occipitotemporal cortex. Neuroimage, 56, 2183–2199. Retrieved from http://www.ncbi.nlm.nih.gov/entrez/query.fcgi?cmd=Retrieve&db=PubMed&dopt=Citation&list_uids=21439386

Yeatman, J. D., Dougherty, R. F., Ben-Shachar, M., & Wandell, B. A. (2012). Development of white matter and reading skills. Proceedings of the National Academy of Sciences of the United States of America, 109(44), E3045–53. https://doi.org/10.1073/pnas.1206792109

